# The role of the LysR-type transcription factor PacR in regulating nitrogen metabolism in *Anabaena* sp. PCC7120

**DOI:** 10.1101/2024.11.01.621236

**Authors:** E. Werner, T. Huokko, A. Santana-Sanchez, S. Picossi, L. Nikkanen, A. Herrero, Y. Allahverdiyeva

**Affiliations:** Molecular Plant Biology, Department of Life Technologies, University of Turku, Turku FI- 20014, Finland; Instituto de Bioquímica Vegetal y Fotosíntesis, Consejo Superior de Investigaciones Científicas, Universidad de Sevilla, Seville E-41092, Spain

**Author notes:** **Correspondance:** Prof. Yagut Allhaverdiyeva, Molecular Plant Biology, Department of Life Technologies, University of Turku, Turku FI- 20014, Finland.

## Abstract

In the filamentous model cyanobacterium *Anabaena* sp. PCC 7120 heterocyst formation is triggered by changes in the C/N-ratio and relies on transcriptional reprogramming in cells. The transcription factor PacR is thought to serve as a global regulator of carbon assimilation under photoautotrophic conditions. In response to C_i_-availability, PacR may modulate the carbon concentrating mechanism and photosynthesis, balancing reducing power generation while protecting the photosynthetic apparatus from oxidative damage. However, PacR also binds to promoters of genes associated with heterocyst formation, although the underlying mechanisms remain unclear. To explore this, we studied a response of a PacR-deletion mutant to a nitrogen source shift from ammonium to nitrate. The absence of PacR led to the heterocyst formation in nitrate containing media, as well as reduced growth and chlorophyll-content. We observed impaired nitrate uptake and disrupted ammonium assimilation via the GOGAT-cycle. This phenotype may be exacerbated by reduced PSI-yield and reduced expression of ferredoxin, which may lead to less reducing equivalents for nitrogen assimilation.

Our results provide insights into the regulation of heterocyst-formation in *Anabaena*, potentially advancing its use in biotechnological applications that utilize heterocyst as microoxic cell factories for N2-fixation and hydrogen production.

## Introduction

The filamentous cyanobacterium *Anabaena* sp. PCC7120 (hereafter *Anabaena*) is a model heterocyst-forming strain that carries out oxygenic photosynthesis and CO_2_-fixation in vegetative (photosynthetic) cells, while performing nitrogen fixation in specialized cells called heterocysts. In the absence of combined nitrogen about 10% of the vegetative cells differentiate into intercalary heterocysts within 24h (Zeng & Zhang, 2022), which form at intervals of 10-15 vegetative cells along the filament (Flores et al., 2019; Nieves-Morión et al., 2021). Heterocysts are specialized microoxic cells that fix atmospheric N_2_ into NH_3_, a process catalyzed by the heterocyst-specific O_2_-sensitive nitrogenase enzyme (Flores et al., 2019; Nieves-Morión et al., 2021). In the heterocyst, microoxic conditions are maintained by high respiration rates (Flores et al., 2019; Harish & Seth, 2020), the flavodiiron (Flv) 3B homo-oligomer catalyzing O_2_ photoreduction (Ermakova et al., 2014), a non-oxygen-evolving Photosystem (PS) II with a reduced antenna size (Ferimazova et al., 2013; Harish & Seth, 2020; Magnuson, 2019) and the formation of a heterocyst envelope consisting of an inner glycolipid (Hgl) layer and an outer polysaccharide (Hep) layer (Cumino et al., 2007; Flores et al., 2019). Vegetative cells supply heterocysts with carbon in the form of sucrose, alanine and glutamate (Burnat et al., 2014). In heterocysts, the NH_4_^+^ produced during N_2_ fixation is incorporated into glutamate by glutamine synthetase, producing glutamine, which is then transferred to the vegetative cell along with beta-aspartyl-arginine, the degradation product of cyanophycine (Flores et al., 2019). The process of heterocyst differentiation is highly complex and tightly regulated, with the primary factor being the C/N balance. In *Anabaena*, the primary trigger for differentiation are rising levels of 2-oxoglutarate (2-OG), which serves as an indicator of the C/N balance. 2-OG is linking carbon and nitrogen metabolism through NH_4_^+^-fixation onto glutamate, which is synthesized from 2-OG, forming glutamine (Zeng & Zhang, 2022; Zhang et al., 2018). The global transcription factor NtcA binds 2-OG and activates the expression of the master regulator of heterocyst formation, HetR, via the response regulator-like factor NtcA (Flores et al., 2019; Zeng & Zhang, 2022; Zhang et al., 2018). NtcA and HetR together orchestrate the developmental process of heterocyst formation (Flores et al., 2019; Zeng & Zhang, 2022). Early in this process, PatS and PatX help to establish the initial heterocyst pattern (Flores et al., 2019). The DevH transcriptional factor regulates heterocyst-specific genes at later stages of differentiation and plays a central role in regulating Hgl layer formation as well as expression of the *cox2* and *cox3* operons (functioning in O_2_ reduction) and the *nif* gene cluster (for N_2_-fixation) (Zeng & Zhang, 2022). Other late-stage regulators involved in heterocyst formation include the NtcA co-activator PipX and HetN, which is crucial for pattern maintenance (Flores et al., 2019).

LysR-type transcriptional regulators (LTTRs) are the largest known family of bacterial DNA-binding proteins, playing a diverse role in regulating a wide range of biological processes. They may act as activators or repressors and often in response to a co-inducer molecule (López-Igual et al., 2012). Among the LTTRs subfamily, the CbbR factors specifically control genes involved in the Calvin-Benson-Bassham (CBB) cycle. In *Anabaena*, there are three CbbR-like LTTRs. One of them, CmpR (*all0862*), activates the *cmp* operon (*alr2877-alr2880*), which encodes an ABC transporter complex for HCO_3_^-^ transport. CmpR also controls its own expression in response to C_i-_ limitation and coordinates this with nitrogen availability through its response to NtcA (Picossi et al., 2015). Another member, CcmR (also known as NdhR), acts as a repressor for carbon transporters and its own expression (López-Igual et al., 2012; Picossi et al., 2015).

The third member, PacR (*rbcR*), is a global regulator. In *Anabaena*, PacR regulates a large number of genes associated with C_i_-fixation (e.g. elements of the carbon concentrating mechanisms (CCM) and RuBisCo). Remarkably, PacR was the first transcriptional factor described in cyanobacteria that can adjust the expression of photosynthetic genes (e.g. the core PSI reaction center *psaA* gene) to C_i_-availability, with most genes showing upregulation by PacR under C_i_-limitation (Picossi et al., 2015). Noteworthy, most photosynthetic genes targeted by PacR probably function in protection against reactive oxygen species, which can be generated due to C_i_-limitation or exposure to high light. It has been proposed that PacR plays a crucial role in modulating the generation of reductive power based on C_i_-availability while safeguarding the photosynthetic apparatus from oxidative stress. Notably, the PacR expression appears to be independent of the C/N regime (Picossi et al., 2015). Despite PacR being identified as a global transcription factor in *Anabaena*, its role has primarily been studied in relation to the regulation of C_i_-assimilation and photosynthetic genes, while its potential role in other parts of the global metabolic network, including nitrogen assimilation, remains largely unexplored. In this study, we analyzed a PacR-deletion mutant using transcriptomics, combined with metabolic and physiological measurements, including the monitoring of photosynthetic capacity, NO_3_^-^ as available nitrogen source in the growth medium. Our results suggest that PacR plays a role in regulating nitrogen metabolism including adaptation to NO_3_^-^ as a nitrogen source, thus influencing heterocyst differentiation in NO_3_^-^ medium. This study broadens the understanding of the PacR regulon, highlighting its role in facilitating adaptation to various nitrogen sources and contributing to heterocyst formation alongside other key transcriptional regulators.

## Materials and Methods

### Strains and culture conditions

The Δ*pacR* deletion mutant of the *Anabaena* sp. PCC7120 wild type strain (CSS74) and control strain with a spectinomycin resistance cassette (CSS77) were described previously (Picossi et al., 2015). The starting cultures were maintained in BG11 medium buffered with 20 mM HEPES-NaOH (pH 7.5) and supplemented with 25 ug/ml of spectinomycin. The cultures were grown at 30°C with 1% CO_2_, ∼50 µmol photons m_-2_ s_-1_, and agitated at 80 rpm. Pre-cultures were generated by harvesting cells from starting cultures after 4 days and resuspending cells at OD_750_=0.1. Pre-cultures were then grown for 4 days in BG11_0_ medium buffered with 20 mM HEPES-NaOH (pH 7.5) supplemented with 6mM NH_4_Cl on day 0, and additional 3mM NH_4_Cl were supplemented on days 2 and 3. Experimental cultures were generated similar to pre-cultures and grown in BG11_0_ medium (pH 7.5) supplemented with either 3 mM NH_4_^+^ or 17.6 mM NO_3_^-^ as the nitrogen source.

### Chl *a* determination and heterocyst frequency

Chl *a* was extracted from cells with 90% methanol and the concentration was determined by measuring OD_665_ and multiplying it with the extinction coefficient factor 12.7 (Meeks & Castenholz, 1971). For counting of heterocysts, Alcian Blue was used to stain the polysaccharide layer of the heterocyst envelope as previously described (Santana-Sánchez et al., 2023). Stained samples were visualized using a Wetzlar light microscope (Leitz, Wetzlar, Germany) and x400 magnification micrographs were taken via mounted camera (Leica). Images were processed using the Leica Application Suite (Leica Microsystems). For each sample, 1000–2000 cells were counted.

### Photosynthetic activity measurements

#### Chl fluorescence and absorbance

A pulse amplitude modulated fluorometer Dual-PAM-100 (Walz) was used to simultaneously monitor Chl *a* fluorescence and P700 absorbance. Cells were harvested and resuspended in fresh BG11 medium to a Chl *a* concentration of 15 µg ml^-1^ and acclimated 1 h under growth conditions. Prior to measurements the samples were dark-adapted for 10min. The effective yield of PSII [Y(II)] and PSI [Y(I)], as well as the acceptor side limitation of PSI [Y(NA)] and donor side limitation [Y(ND)], were determined as described previously (Huokko et al., 2017; Schreiber et al., 2008). Actinic light (630 nm, 50 µmol photons m^−2^ s^−1^), saturating pulses (300 ms, 5,000 μmol photons m^-2^s^-1^), and strong far-red light (720 nm, 75 W m^−2^) were applied during analysis.

#### Membrane Inlet Mass Spectrometry

*In vivo* measurements of ^16^O_2_ (m/z=32), ^18^O_2_ (m/z=36) and CO_2_ (m/z=44) fluxes were monitored using Membrane Inlet Mass Spectrometry (MIMS) as described previously (Santana-Sánchez et al., 2023). Cells were harvested after 48h and resuspended in fresh BG11 medium, adjusted to Chl *a* 10 µg ml^-1^ and acclimated for 1 h under growth conditions prior to measurements. For all MIMS measurements, gas exchange was monitored for 4min in darkness, followed by 5min of high light illumination (500 µmol photons m^-2^ s^-1^) and for an additional 2min in the dark.

#### Determination of nitrogen uptake and intracellular carbon and nitrogen content

NO_3_^-^ concentration in the growth media was determined spectrophotometrically (Rice et al., 2012). To determine intracellular C/N ratio, cells were harvested 48h after the shift to NO_3_^-^ as nitrogen source and 10mg of biomass was lyophilised. Total carbon and nitrogen analysis was conducted using a FLASH 2000 Organic Elemental Analyzer (Thermo Scientific).

### Metabolite extraction and quantification

Cells were harvested by filtration on Millipore HATF nitrocellulose membranes (0.45 μm) from cultures grown for 4 days with NH_4_^+^ as the nitrogen source, as well as at 1 h and 12 h after shifting to NO_3_^-^ containing growth medium. Filters were first frozen in liquid N_2_, then harvested cells were resuspended in PBS buffer, pelleted and stored at -80°C. Metabolites were extracted by adding 400 μL of cold extraction solvent (acetonitrile:methanol:H_2_0; 40:40:20) to each sample, followed by three cycles of sonication (60sec, Elma Elmasonic P) and vortexing (120 sec). The samples were then centrifuged at 14 000rpm for 20 min at +4°C, after which the supernatants (350 µl) were transferred to evaporation tubes and dried under a N_2_-stream. Dried samples were resuspended in 50 µl cold extraction solvent and transferred to HPLC auto sampler glass vials. The analysis was performed by injecting 2μL of sample extract to a Thermo Vanquish UHPLC coupled with a Q-Exactive Orbitrap quadrupole mass spectrometer equipped with a heated electrospray ionization (H-ESI) source probe (Thermo Fischer Scientific). Chromatographic separation of metabolites was done using a SeQuant ZIC-pHILIC (2.1 × 100mm, 5μm particle) column (Merck), which was maintained at +40°C. Scanning was performed in mass range 55 to 825m/z. Gradient elution was carried out with acetonitrile and 20mM ammonium hydrogen carbonate in water, adjusted to pH 9.4, as mobile phases. TraceFinder 4.1 software (Thermo Fischer Scientific) was used for data analysis.

### RNA isolation and sequencing

Cells were harvested by filtration on Millipore HATF nitrocellulose membranes (0.45μm) from cultures grown for 4 days with NH_4_^+^ as the nitrogen source, as well as at 1 h and 12 h after shifting to NO_3_^-^ containing growth medium. Cells were resuspended in RNA-resuspension buffer (Walter et al., 2016), centrifuged for 3min at 4°C and then the pellet was then frozen in liquid N_2_. Total RNA was isolated using the hot-phenol method (Walter et al., 2016), with the incubation step in phenol:chloroform:iso-amyl alcohol performed at 65°C instead of 95°C. Finally, the samples were resuspended in 25 µl of Milli-Q Water. DNAse treatment was performed with the invitrogen kit. RNA concentration was measured with DS-11+ spectrophotometer (DeNovix) and 3ug of RNA from each sample was sent to BGI (China) for RNA-sequencing. The RNA was sequenced using DNBseq with a read length of PE 100bp and data was analyzed with the Chipster software. Initial trimming of the reads was performed using FastX, followed by alignment with the genome of *Anabaena* (downloaded from EnsemblBacteria (bacteria.ensembl.org), nostoc_sp_pcc_7120_fachb_418_gca_000009705, version ASM970v1_, current ENSEMBL release, October 2023) using TopHat2. Aligned reads per gene were then counted with HTSeq. Differential expression analysis was conducted using DESeq2 (*log*_*2*_*FC* ≧ I1I, p<0.05). For experimental quality control, PCA analysis was performed (Suppl. Fig. 5).

## Results

### Phenotype of the Δ*pacR* mutant under different nitrogen sources

The growth phenotypes of the Δ*pacR* mutant and the control strain (CS) were compared under different nitrogen replete conditions. The Δ*pacR* mutant demonstrated significantly slower growth, reaching an optical density (OD_750_) of only 72.3±11.15% of that of the CS when NH_4_^+^ was used as the nitrogen source in the medium (Suppl. Fig. 1). When grown with NO_3_^-^ as the only nitrogen source in the medium, the Δ*pacR* mutant reached only 80.4±16.5% of the OD_750_ of the CS (Fig.1 A), along with a reduction to only 64.2±16.9% of the Chl-content of the CS (Fig. 1 B) and only 78.4±15.6% of the Chl/OD ratio of the CS (see Fig. 1 C). Although both the Δ*pacR* mutant and the CS exhibited almost no heterocyst differentiation in NH_4_^+^-containing medium (the frequency of heterocysts was 0.05±0.04% of total cells for the CS and 0.26±0.36% for Δ*pacR* (Suppl. Table 1)), the Δ*pacR* mutant showed a significantly higher proportion of heterocysts (7.8±1.4%, p<0.05) compared to the CS (0.6±0.5%) when cultured in NO_3_^-^-containing medium (Fig. 1 D; Suppl. Table 1 and Suppl. Fig. 2). These results suggest that PacR is required for the suppression of heterocyst differentiation in the presence of NO_3_^-^.

**Figure 1.**
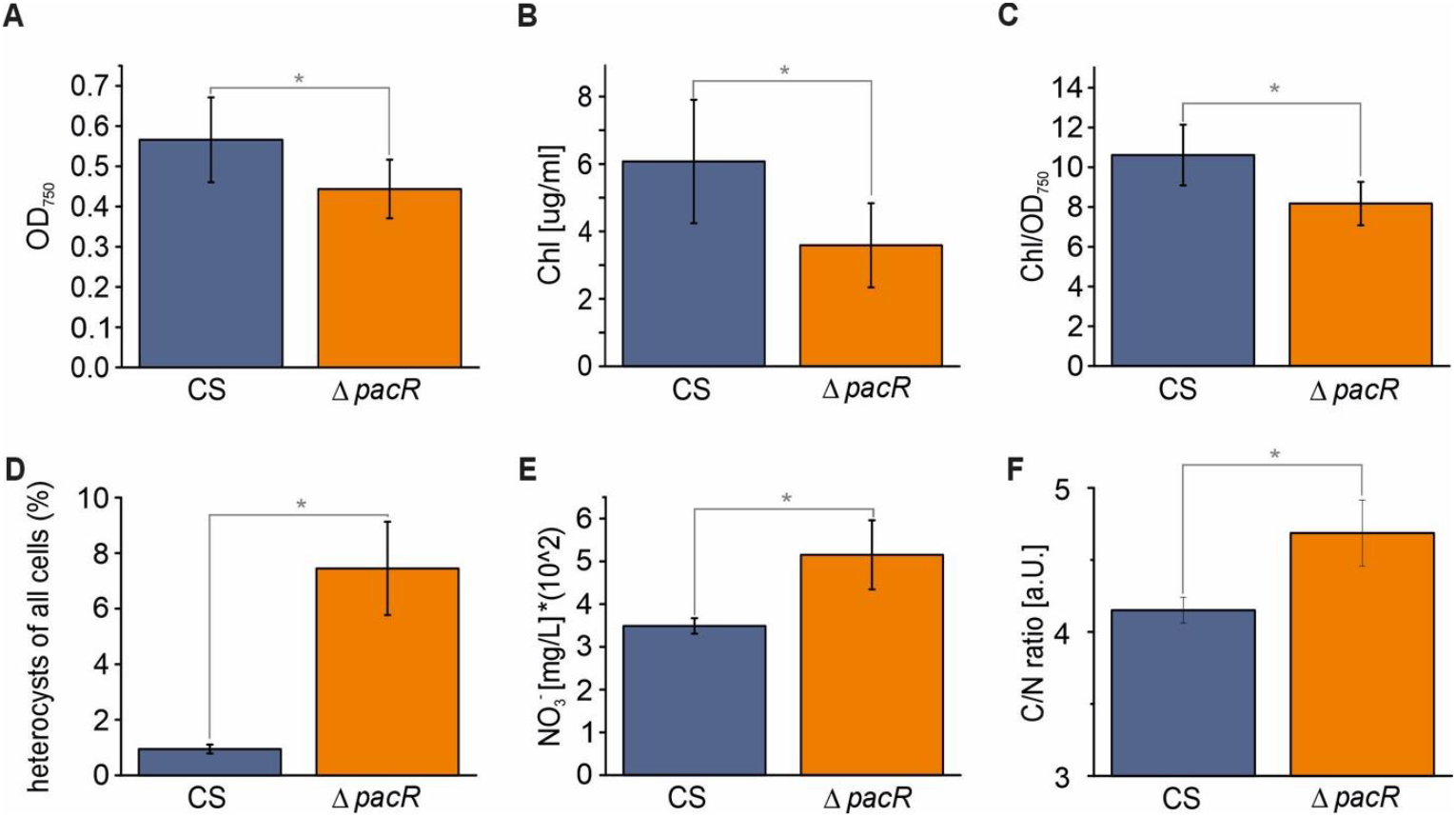
Phenotypic characteristics of the Δ*pacR* mutant and the control (CS) strain 48 h after the shift from NH_4_^+^ to NO_3_^-^. **A**. OD_750_, **B**. Chl content. **C**. Chl/OD ratio. **D**. number of heterocysts. **E**. NO_3_^-^ in the growth medium. **F**. C/N ratio measured by elemental analysis. Values are mean ± SD, n=3 biological replicates. In D. 1000-2000 cells were counted for each biological replicate. Asterisks indicate statistically significant differences (t-test, p<0.05) compared to the CS.

Next, we choose to study the Δ*pacR* mutant during the transition from NH_4_^+^ to NO_3_^-^ as the nitrogen source, focusing on the potential mechanisms driving increased heterocyst differentiation in the presence of NO_3_^-^. The *ΔpacR* mutant demonstrated impaired NO_3_^-^ uptake during the 48 h after transition, showing 1.5±0.2 times more residual NO_3_^-^ in the growth medium compared to the CS (Fig. 1 E). Importantly, organic elemental analysis of the cells showed an increased C/N ratio in the Δ*pacR* mutant, with a 14.3±4.9% reduction in total N content compared to CS (Fig. 1 F; Suppl. Table 2), indicating a greater deprivation in N relative to C.

### Photosynthetic characterization of the Δ*pacR* mutant

Next, we examined the photosynthetic capacity of the Δ*pacR* mutant cultured with NO_3_^-^. The effective yield of PSI, Y(I), was reduced in the mutant compared to the CS (Fig. 2 A) and acceptor-side limitation of PSI, Y(NA), was increased (Fig. 2 B). Although the effective PSII yield, Y(II), was higher in the mutant, this difference was not statistically significant (Suppl. Fig. 3 A). Additionally, donor side limitation Y(ND) of PSI remained unaffected (Suppl. Fig. 3 B).

**Figure 2.**
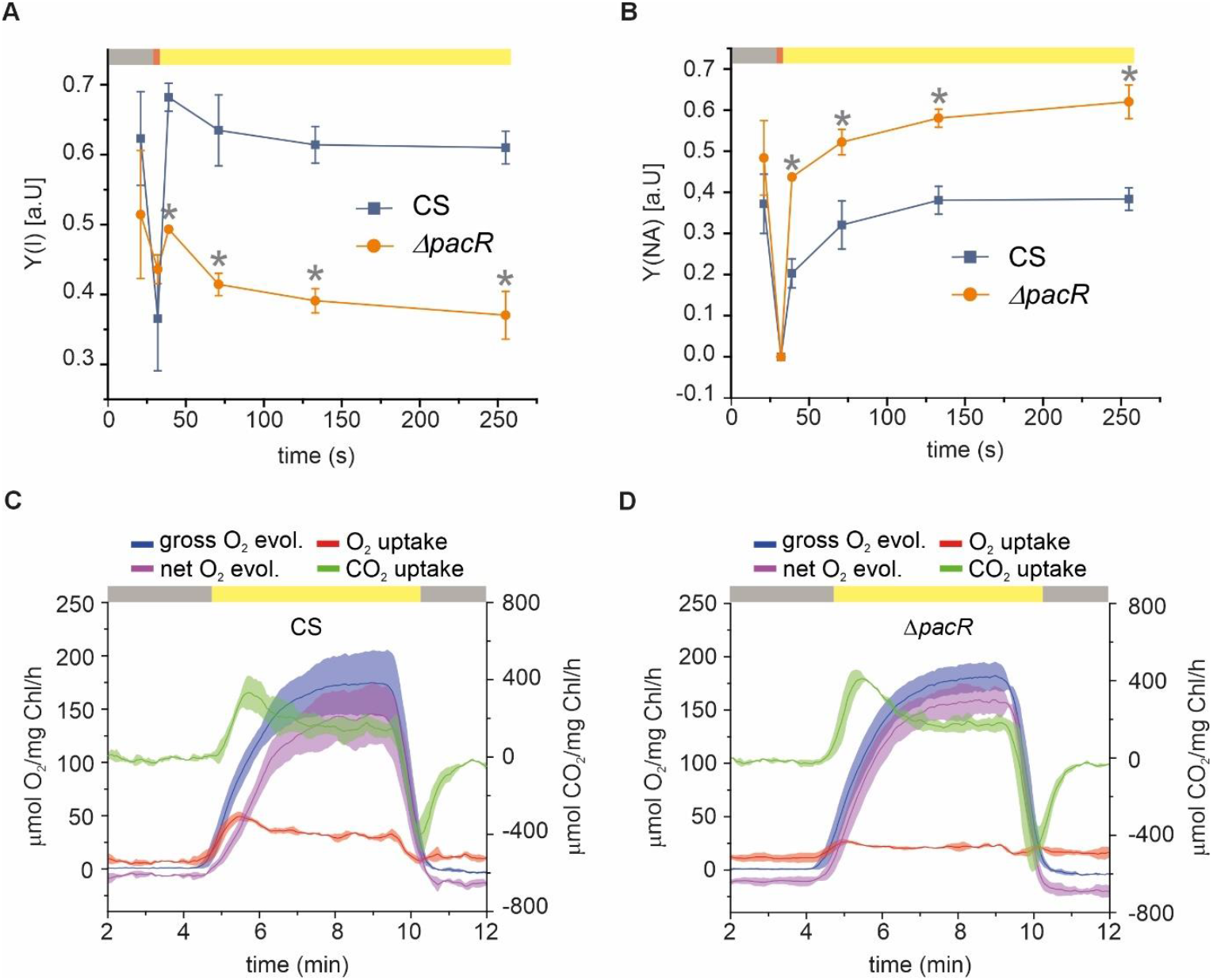
Photosynthetic performance of the *ΔpacR* mutant and the control (CS) strain in the presence of NO_3_^-^. **A**. the effective PSI yield [Y(I)] and **B**. the acceptor side limitation of PSI [Y(NA)] measured in darkness (grey bars), under illumination with far-red (FR) light (red bars) and with 50 µmol photons m-2 s-1 of actinic white light (yellow bars) **C**. and **D**. O_2_ and CO_2_ exchange rates of the CS (C) and the Δ*pacR*-mutant (D). Gas exchange rates were recorded in darkness (gray bars) and under illumination with 500 µmol photons m^-2^ s^-1^ of actinic white light (yellow bars). Values are Mean ± SD, n=3 biological replicates. Asterisks indicate statistically significant differences (t-test, p<0.05) compared to the CS.

For a more detailed assessment of the activity of the photosynthetic apparatus, we monitored real-time gas exchange fluxes via Membrane Inlet Mass Spectrometry (MIMS) and used the ^18^O_2_ isotope to differentiate oxygen uptake from evolution. The illumination of the CS led to rapid O_2_-photoreduction, primarily attributed to Flvs (Santana-Sánchez et al., 2023), reaching a maximum of 40.9±3.9 μmol mg Chl a^-1^ h^-1^ (p<0.05)) and plateaued at 23.1±2.7 μmol mg Chl a^-1^ h^-1^ (p<0.05). However, the Δ*pacR* mutant exhibited significantly lower O_2_-photoreduction, reaching a maximum of 13.0±3.0 μmol mg Chl a^-1^ h-1, with an average plateau level of 7.3±5.0 μmol mg Chl a^-1^ h^-1^ (p<0.05), indicating impaired light-induced O_2_-uptake in the Δ*pacR* mutant. This is in line with the higher net O_2_ evolution observed in the mutant compared to the CS, calculated as the difference between gross O_2_ production and O_2_ photoreduction, with values of 142.1±26.1 and 159.4±11.1 μmol mg Chl a^-1^ h^-1^ for the CS and Δ*pacR*, respectively. Gross photosynthetic O_2_ evolution exhibited no significant difference between the CS and Δ*pacR*. Additionally, dark respiration as well as maximal and steady-state CO_2_ uptake were somewhat higher in the mutant (Fig. 2 C, D; Supp. Table 3).

### Metabolic analysis of the Δ*pacR* mutant

To understand the metabolic responses associated with heterocyst formation in the Δ*pacR* mutant cultured with NO_3_^-^, we monitored the levels of several key metabolites from the Citric Acid (TCA)-, Glutamine Oxoglutarate Aminotransferase (GOGAT)- and Ornithine-Ammonia (OAC)-cycles at three time points: in the presence of NH_4_^+^ as the only supplemented nitrogen source in the medium just before the shift (t=0 h), and 1 h and 12 h after the shift to NO_3_^-^ containing medium. Changes in metabolite levels in the Δ*pacR* mutant in relation to the CS are shown for each time point (Fig. 3 A-C), as well as changes of metabolite levels in relation to t=0 h for each strain (Supp. Fig. 4 A, B). Differences in metabolite levels between these two strains were exacerbated with duration of exposure to NO_3_^-^. Most metabolites analyzed showed lower levels in Δ*pacR* compared to the CS, except for alanine (Fig. 3 A-C). Upon shifting to NO_3_^-^, levels of 2-OG increased in both strains in relation to t=0 h (Supp. Fig. 4 A, B), with a statistically significant increase observed only in the CS. Alanine increased significantly in the Δ*pacR* mutant in relation to the CS at t=12 h (*log*_*2*_*FC=*1.74 ±0.69) (Fig. 3 C). Strikingly, glutamate levels were lower in Δ*pacR* in relation to the CS at all timepoints (for t=1 h the *log*_*2*_*FC=* -0.44±0.18 and for t=12 h the *log*_*2*_*FC=* -0.93±0.31) (Fig. 3 A-C), while glutamine showed the same trend. Interestingly, aspartate, arginine and ornithine, which are intermediates of the OAC and cyanophycine synthesis, were found in lower amounts in the *ΔpacR* mutant at all timepoints, while asparagine levels were higher at all timepoints (Fig. 3 A-C). Pyruvate content peaked 1 h after the shift in both strains (Supp. Fig. 4 A-B), with lower pyruvate levels in Δ*pacR* compared to the CS at all timepoints (Fig. 3 A-C). Similarly, citric acid, fumarate and malate, intermediates of the TCA cycle, were less abundant in Δ*pacR* compared to the CS (Fig. 3 A-C).

**Figure 3.**
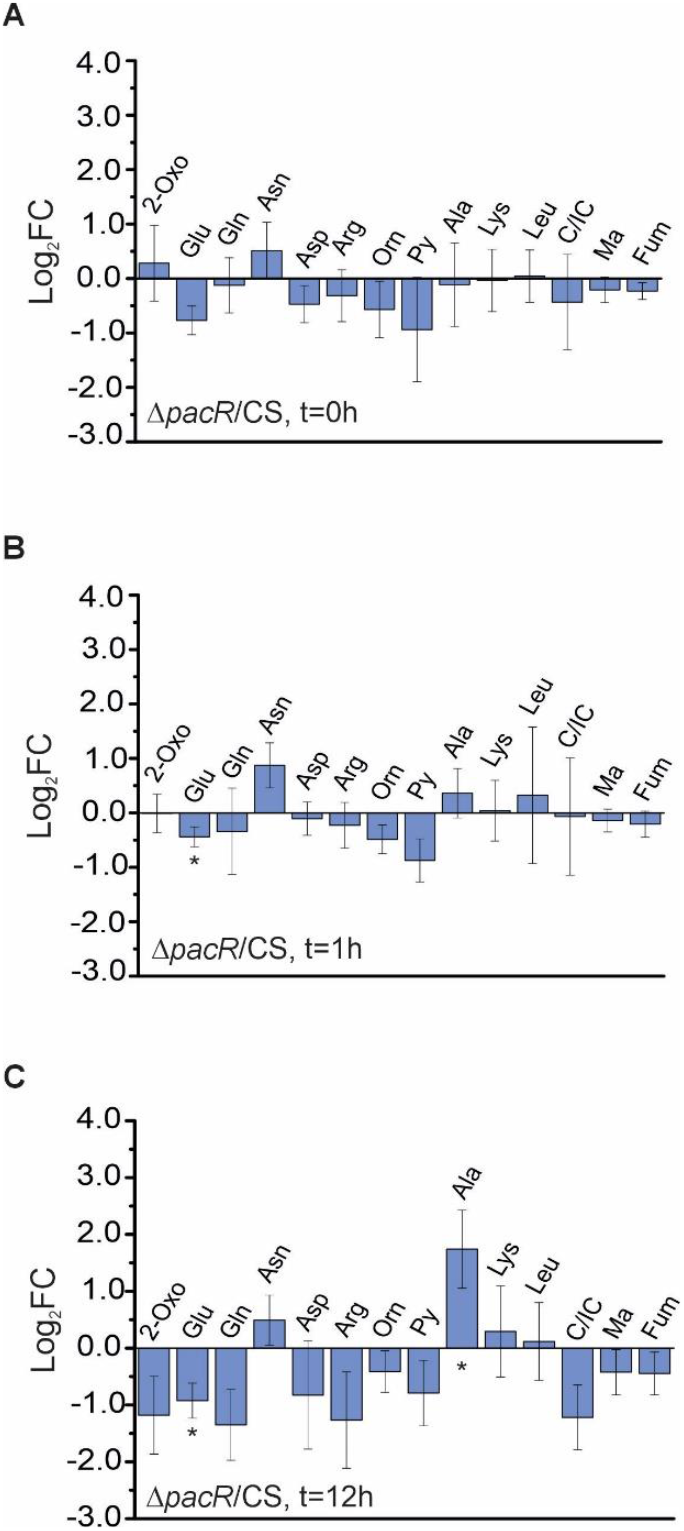
Metabolic analysis of the *ΔpacR* mutant before and at 1 h and 12 h after the shift from NH_4_^+^ to NO_3_^-^ as the only supplemented nitrogen source in the medium. **A**. metabolite levels of *ΔpacR* relative to the control strain (CS) at t=0 h **B**. metabolite levels of *ΔpacR* relative to CS at t=1 h. **C**. metabolite levels of *ΔpacR* relative to CS at t=12 h. 2-OG=2-Oxoglutarate, Glu=Glutamate, Gln=Glutamine, Asn=Asparagine, Asp=Aspartic acid, Arg=Arginine, Orn=Ornithine, Py=Pyruvate, Ala=Alanine, Lys=Lysine, Leu=Leucine, CI/C=Citrate/Isocitrate, Ma=Malic acid, Fum=Fumarate. Values are mean±SD (n=3-6 biological replicates) and shown as *log*_*2*_*FC*. Asterisks indicate statistically significant differences, calculated based on the normalised total ion count (t-test, p < 0.05).

### RNA-seq analysis of the Δ*pacR* mutant

To elucidate the differential expression of genes due to the deletion of *pacR*, we performed RNA-seq analysis of the CS and the *ΔpacR* mutant grown in the presence of NH_4_^+^ as the only supplemented nitrogen source in the medium and sampled just before the shift (t=0 h) as well as 1 h and 12 h after the shift to medium supplemented with NO_3_^-^ as the only nitrogen source. At t=0 h, 705 genes were found to be upregulated and 774 genes downregulated, at t=1 h, 639 genes were upregulated and 707 genes downregulated and at t=12 h, 1344 genes were upregulated and 1034 genes downregulated in the *ΔpacR* mutant in relation to the CS. Thus, up to ∼38% of the gene transcripts of *Anabaena* (RefSeq: GCF_000009705.1) were differentially expressed in the Δ*pacR* mutant at any given time. Differentially expressed genes were found e.g. in the categories of global transcriptional regulators, nitrogen uptake and heterocyst formation, the GOGAT-cycle, the Ornithine-Ammonia Cycle and cyanophycine synthesis, carbon uptake as well as the photosynthetic machinery.

#### Transcriptional regulators

The transcription level of PacR (*rbcR*) in the CS remained independent of the nitrogen source, showing no differential expression between timepoints (Fig. 4; Suppl. Tables 7,8), as reported previously (Picossi et al., 2015). Strikingly, NtcA (*ntcA*) transcripts were slightly but significantly upregulated at t=12 h in the *ΔpacR* compared to the CS (Fig. 4; Suppl. Table 6). Additionally, the transcript levels of another global transcriptional regulator, FurA (*furA*), were slightly elevated in the mutant 1 h after the shift, as the transient downregulation observed in the CS was absent in the mutant (Fig. 4; Suppl. Tables 5,7,8). Moreover, no differential expression of CmpR-transcripts was observed between the CS and the Δ*pacR* mutant.

**Figure 4.**
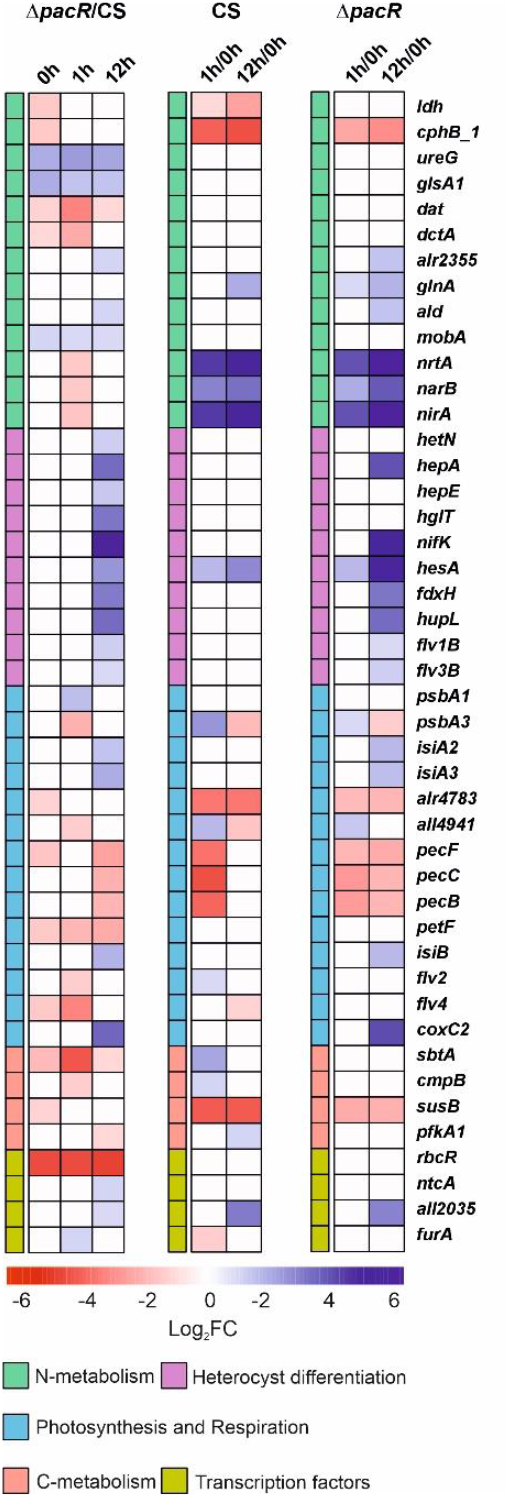
Expression patterns in the control strain (CS) and the Δ*pacR* mutant during the shift from NH_4_^+^ to NO_3_^-^ as only nitrogen source in the medium. **Δ*pacR*/CS**. Differential gene expression between the **Δ***pacR* and CS *a* before, at 1 h and 12 h after the shift to NO_3_^-^. **CS-column**. Differential gene expression in the CS between t=0 h and t=1 h and between t=0 h and t=12 h. **Δ*pacR*-column**. Differential gene expression in **Δ***pacR* between t=0 h and t=1 h and between t=0 h and t=12 h (log_2_FC ≧ I1I, p<0.05).

#### Nitrogen metabolism

The NirA-operon (*nirA*-*nrtABCD*-*narB*), which codes for the NO_3_^-^ uptake machinery (Frías & Flores, 2010), was highly upregulated in the CS 1 h after the shift to NO_3_-whereas this upregulation was strikingly delayed in the Δ*pacR* mutant, with a slight but significantly lower transcript level at t=1 h (Fig. 4; Suppl. Tables 5,7,8).

Related to the GOGAT-cycle, the transcript levels of glutaminase 1 (*glsA1*) (Zhou et al., 2008) was slightly upregulated in Δ*pacR* compared to the CS at all time points (Fig. 4; Suppl. Tables 4-6), while the transcript of glutamine synthetase (*glnA*) (Álvarez-Escribano et al., 2024) was slightly upregulated in Δ*pacR* 1 h after the shift and in both strains 12 h after the shift (Fig. 4; Suppl. Tables 7,8). The transcript levels of *all0396*, whose promoter is directly bound by PacR (Picossi et al., 2015), were downregulated in Δ*pacR* compared to the CS, showing slight downregulation at both t=0 h and t=12 h and intermediate downregulation at t=1 h (Fig. 4; Suppl. Tables 4-6). L-2,4-diaminobutyrate:2-ketoglutarate 4-aminotransferase (DABA AT) is encoded by *all0396*, and it is involved in the biosynthesis of schizokenin, an iron-chelating siderophore in *Anabaena* (Su et al., 2024; Wellham, 2017). The transcript level of alanine dehydrogenase (*ald*) (Pernil et al., 2010) was slightly upregulated in *ΔpacR* compared to the CS at t=12 h (Fig. 4; Suppl. Tables 6-8).

#### Heterocyst formation

Many genes associated with heterocyst formation were upregulated in the *ΔpacR* strain 12 h after the shift compared to the CS. Unsurprisingly, NtcA (*ntcA*) showed slightly increased expression (Fig. 4; Suppl. Table 6). Genes involved in the formation of the Hep-layer, such as *hepA* (X. P. Wang et al., 2016; Y. Wang et al., 2007), as well as various glycosyl transferases, like *hepE* (X. P. Wang et al., 2016) were upregulated in various degrees. A glycolipid synthase (*hglT*) involved in Hgl-layer formation (Zeng & Zhang, 2022), was also intermediately upregulated. Several nitrogenase-related genes were upregulated, including intermediate to strong upregulation of the main *nif*-gene cluster as well as intermediate upregulation of *nifV*, belonging to the additional *nif*-gene cluster found in *Anabaena* (Stricker et al., 1997). Concomitantly, the heterocyst-specific ferredoxin (*fdxB*) (Pils et al., 2004) and the heterocyst-specific uptake hydrogenase genes (*hupL, hupS)* (Santana-Sánchez, 2021) also showed intermediately increased expression. The heterocyst-specific *flv1B* and *flv3B* transcripts (Ermakova et al., 2013) were slightly upregulated (Fig. 4; Suppl. Tables 6,8).

#### Carbon metabolism

The gene encoding a putative SbtA carbon transporter orthologue (*sbtA*) (Herrero & Flores, 2019), was slightly downregulated in *ΔpacR* compared to the CS and did not undergo transient upregulation at t=1 h as in the CS, resulting in strong downregulation compared to the CS at this timepoint (Fig. 4; Suppl. Tables 4-7). A similar pattern was observed for transcripts of *cmpB*, corresponding to a bicarbonate transporter (Herrero & Flores, 2019), however at t=1 h the gene was only slightly lower expressed than in the CS (a similar tendency was observed for the entire *cmpABCD* operon) (Fig. 4; Suppl. Tables 5,7). Moreover, sucrose-synthase (*susB*) transcription (Katoh et al., 2004) was slightly downregulated at t=0 h in Δ*pacR* compared to the CS whereas in both strains the gene was downregulated after the shift, albeit only intermediately in the mutant and strongly in the CS (Fig. 4; Suppl. Tables 4,7,8).

#### Photosynthesis and respiration

Interestingly, the transcription of PSII core protein D1 (*psbA1*) (Sicora et al., 2009) was slightly upregulated 1 h after the shift in the mutant in relation to the CS (Fig. 4; Suppl. Table 5). In contrast, the transient upregulation at t=1 h of *psbA3* (Summerfield et al., 2008) was only minor in *ΔpacR* while it was moderate in the CS (Fig. 4; Suppl. Tables 5,7,8). Transcripts of *isiA2* and *isiA3*, iron-stress-induced proteins with homology to CP43 (Nagao et al., 2021), were slightly upregulated after 12 h in Δ*pacR* in relation to the CS (Fig. 4; Suppl. Tables 6,8).

Before the shift, the phycoerythrocyanin (*pec)* operon, which encodes the components of the light-harvesting phycobilisome (PBS) complex (Gollan et al., 2020), was slightly downregulated in Δ*pacR* compared to the CS (significantly for *pecE* and *pecF*). Genes of this operon were then downregulated to about the same level in both strains at t=1 h and were then significantly expressed at slightly to intermediately higher levels in the CS compared to *ΔpacR* after 12 h (Fig. 4; Suppl. Tables 4,7,8). Transcripts of *all4940* and *alr4783* associated with the orange carotenoid protein (OCP) (López-Igual et al., 2016) were slightly downregulated in Δ*pacR* compared to the CS at t=0 h and then downregulated significantly to similar levels in both strains after the shift (Fig. 4; Suppl. Tables 4,7,8).

The following transcripts of cytochrome c oxidase subunits were moderately to highly upregulated at t=12 h in Δ*pacR* compared to the CS: *coxA2* and *coxC2* corresponding to COX2 (homologous to aa3-type cytochrome c oxidases) as well as *coxA3*, corresponding to COX3 (homologous to alternative respiratory terminal oxidases) (Fig. 4; Suppl. Tables 6,8). Both COX2 and COX3 were previously found to be upregulated under nitrogen deficiency (Wünschiers et al., 2006). The transcript levels of vegetative cell-specific ferredoxin 1 (*petF*) (Schmitz & Böhme, 1995), were downregulated, from slight to moderate level, at all timepoints in *ΔpacR* compared to the CS (Fig. 4; Suppl. Tables 4-6), possibly due to PacR binding to the *petF* promoter. In the CS, *flv2* (*all4444*) and *flv4* (*all4446*) (Ermakova et al., 2013) were transiently upregulated at t=1 h (significantly for *flv2*). This response was absent in the mutant, where expression levels remained generally low (Fig. 4; Suppl. Tables 5,7). The expression of *flv1A* (*all3891*) and *flv3A* (*all3895*) (Ermakova et al., 2013) did not differ significantly between the mutant and the CS, and their expression kinetics followed the CS with a non-significant transient increase 1 h after the shift.

## Discussion

In this study, we found that the *ΔpacR* mutant of *Anabaena* forms heterocyst in nitrogen replete conditions (Fig. 1), in a manner similar in diazotrophic conditions (Brenes-Álvarez et al., 2019; Flores et al., 2019), following a shift from NH_4_^+^ to NO_3_^-^ as the nitrogen source in the medium (Fig. 1 D, Suppl. Table 1). The upregulation of genes associated with heterocyst formation occurred in the *ΔpacR* mutant in the 12 h interval after the nitrogen source shift (Fig. 4, Suppl. Tables 2-6). These findings suggest that the deletion of PacR does not directly trigger heterocyst formation. Instead, the response seems secondary to the shift from NH_4_^+^ to NO_3_^-^, which likely disrupts the cellular C/N-balance (Zhang et al., 2018). *Anabaena* typically undergoes heterocyst differentiation within the first 24h after N deprivation (Xing et al., 2022). To gain insights into changes leading up to heterocyst formation, we collected metabolic and RNA-seq data at early time points (0 h, 1 h, 12 h). Additionally, physiological measurements were performed 48 h after the shift, when heterocyst formation was complete.

The *ΔpacR* mutant exhibited a significantly elevated C/N ratio (Fig. 1 F), indicating intracellular nitrogen limitation, likely due to impaired NO_3_^-^ uptake (Fig. 1 E), and delayed induction of NO_3_^-^-transporters in the mutant after the shift to NO_3_^-^ containing medium (Fig. 4, 5; Suppl. Tables 5,7,8). Moreover, PacR may influence the amount of reducing power available from photosynthesis by regulating vegetative cell ferredoxin-1 (*petF*) expression (Picossi et al., 2015). The observed downregulation of *petF* in the *ΔpacR* mutant (Fig. 4, 5; Suppl. Tables 4-8) may be detrimental to conversion of nitrate to ammonium because this reaction is catalyzed by reduced ferredoxin-powered nitrate reductase (NarB) and nitrite reductase (NirA) (Watzer et al., 2019). This would result in observed higher residual NO_3_^-^ levels in the medium at 48 h (Fig. 1 E). Similarly, Zhang et al. (2007) reported that *Anabaena* strains with mutated PII protein (S49A) formed heterocysts even in the presence of nitrate, attributing it to partial impaired nitrate uptake (Y. Zhang et al., 2007).

**Figure 5:**
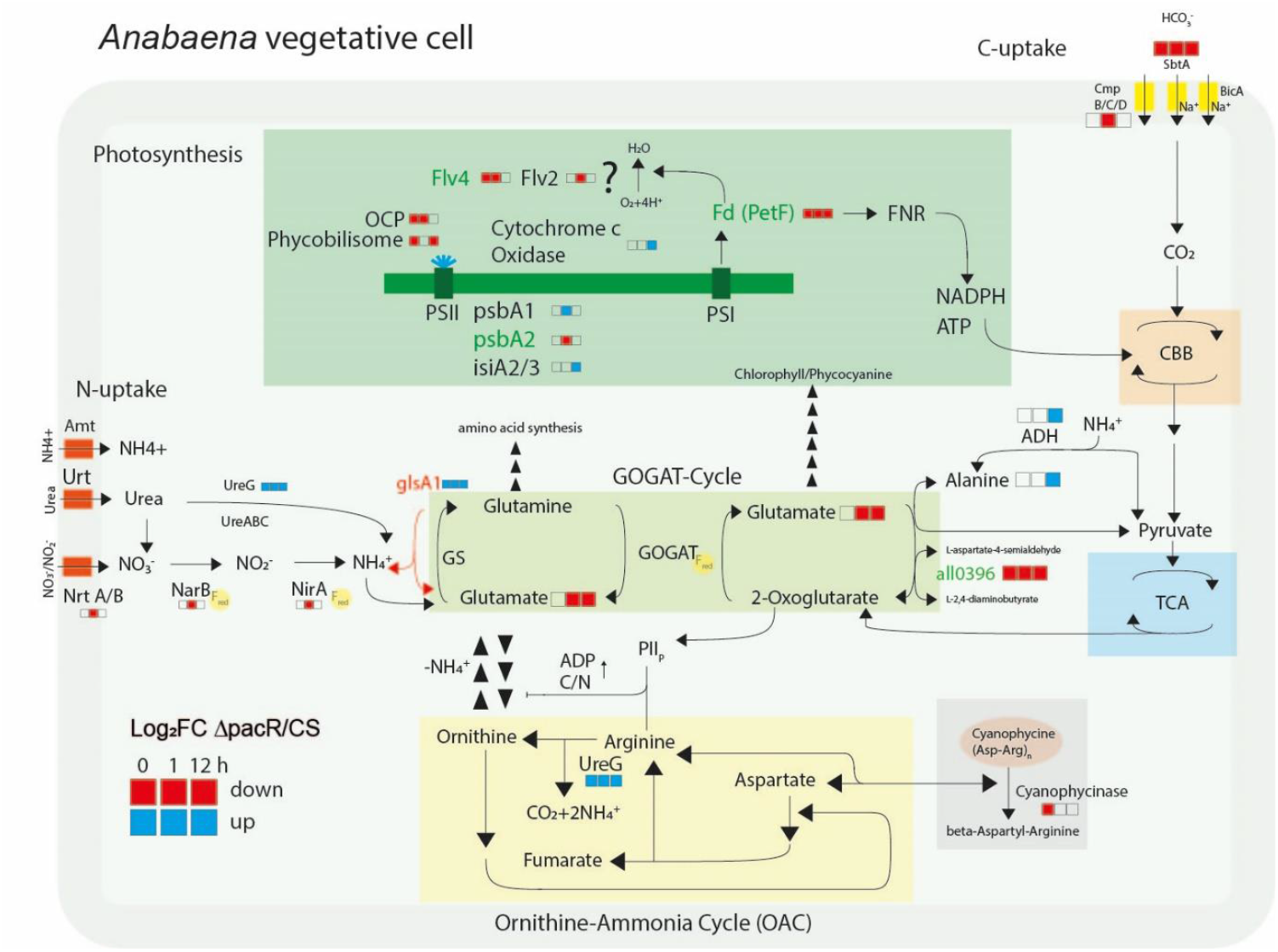
Overview of modified central metabolic pathways in the *ΔpacR* mutant. Shown are pathways involved in C- and N-import, including the GOGAT-cycle, OAC cycle, cyanophycine synthesis, the CBB-cycle and the TCA-cycle. Additionally, the main components of the photosynthetic electron-transport chain are shown. Where reduced ferredoxin is needed for protein function is indicated with a yellow bubble. Boxes indicate genes or metabolites found to be up-(blue) or down-(red) regulated in the *ΔpacR* mutant in relation to the CS before (t=0 h) and 1 h or 12 h after the shift from NH_4_^+^ to NO_3_^-^. Promoters of genes directly bound by PacR (Picossi et al., 2015) are marked in green.

The depletion of free amino acid pools is typical under nitrogen depleted conditions in *Anabaena* (Perin et al., 2021), indicating a lack of nitrogen for metabolic functions. Here, reduced levels of glutamine, glutamate and 2-OG also suggest an imbalance in C/N metabolism. Unlike the usual response in wild-type *Anabaena*, where glutamate levels rise and glutamine levels drop when shifting to diazotrophic conditions (Perin et al., 2021), the Δ*pacR* mutant showed decreases in both glutamate and glutamine, although the changes in glutamine levels were not statistically significant (Fig. 3 A-C, 5). Since glutamate and glutamine are crucial indicators of nitrogen status, this reduction implies a weakened nitrogen assimilation capacity and a reduction in GOGAT-cycle activity. 2-OG, which serves as a signaling molecule for the C/N balance (Robles-Rengel et al., 2019), is known to increase following nitrogen deprivation, triggering heterocyst formation in *Anabaena* (Laurent et al., 2005; Muro-Pastor et al., 2001; Perin et al., 2021; Zeng & Zhang, 2022). However, other factors, like availability of glutamate and glutamine, also play a role in signaling nitrogen status (Forchhammer & Selim, 2020). This is supported by our observation of low 2-OG levels in the Δ*pacR* mutant even though heterocyst formation still occurs, suggesting that the trigger may involve changes in the ratio between 2-OG and other GOGAT intermediates rather than the absolute levels of 2-OG.

The upregulation of the phosphate-dependent glutaminase 1 (*glsA1*) (Fig. 4, 5, Suppl. Tables 4-8), which is mainly responsible for glutamine deamination and is typically downregulated transiently under nitrogen-limiting conditions (Zhou et al., 2008), may also contribute to the disruption of the GOGAT-cycle. Increased Glutaminase activity may disrupt the function of ATP-dependent glutamine synthetase (GS, *glnA*), a critical regulatory point in the GOGAT-cycle that is upregulated when nitrogen is limited (Forchhammer & Selim, 2020; Galmozzi et al., 2010). Notably, inhibiting GS with the specific inhibitor MSX can counteract the repressive effects of fixed nitrogen on nitrogen fixation and heterocyst formation in *Anabaena* sp. (Mishra, 2003).

Alanine levels were significantly higher in the *ΔpacR* mutant compared to the CS following the 12 h shift to NO_3_^-^ (Fig. 3 C, 5). An increase in alanine levels following the shift to diazotrophic conditions was previously observed in *Anabaena* wild type (Perin et al., 2021) and *Anabaena cylindrica* (Rowell & Stewart, 1976), probably because alanine is needed for carbon transfer to heterocysts (Burnat et al., 2014; Pernil et al., 2010). Observed increase of alanine likely derives from the catabolism of nitrogenous compounds, such as cyanophycine degradation (Rowell & Stewart, 1976). Therefore, the elevated alanine levels in this study signify a metabolic state in the *ΔpacR* mutant that resembles diazotrophic conditions, potentially linking them to heterocyst formation. At the same time, the transcript level of alanine dehydrogenase (ADH, *ald*) (Pernil et al., 2010) was slightly upregulated in *ΔpacR* compared to the CS at t=12 h (Fig. 4, 5; Suppl. Tables 6-8). In *Anabaena* spp., ADH is either exclusively expressed or exhibits higher expression levels following the shift to diazotrophic conditions (Pernil et al., 2010). Furthermore, ADH activity has been shown to increase under nitrogen deficiency in *Anabaena cylindrica* (Rowell & Stewart, 1976). These observations indicate that the transcriptomic changes in *ΔpacR* reflect a metabolic adaptation consistent with diazotrophic conditions.

Interestingly, the ADH pathway also possesses synthetic activity, converting pyruvate and NH_4_^+^ into alanine, which facilitates NH_4_^+^-assimilation under conditions of GOGAT-cycle inactivation (Flores & Herrero, 1994; Meeks et al., 1977). In this study, the GOGAT-cycle impairment is likely due to reduced GS efficiency and previous research indicates that NH_4_^+^ incorporation through ADH increases when GS is inactivated by MSX treatment (Meeks et al., 1977). This suggests that there may be increased synthetic ADH activity in the mutant, which may contribute to the increased alanine levels and provide a compensatory mechanism, that serves to counterbalance decreased operation of the GOGAT-cycle.

Moreover, PacR, by regulating SbtA and influencing CmpB expression, may play a crucial role in adjusting carbon intake relative to C/N balance (Fig. 4; Suppl. Tables 4-8), and the observed reduction in their gene expression in the Δ*pacR* mutant could explain its inability to replenish 2-OG levels and the subsequent GOGAT-cycle intermediates.

The reduced PSI yield in the Δ*pacR* mutant is mainly due to increased acceptor-side limitation (Fig. 2 A, B) linked to significantly decreased O_2_ photoreduction (Fig. 2). Since O_2_-photoreduction is mainly attributed to Flv proteins (Santana-Sánchez et al., 2023) the downregulation of *flv2* and *flv4* transcripts in the mutant (Fig. 4, 5; Suppl. Tables 5, 7) along with previous findings that PacR binds to the promoter regions of *flv1A* and *flv4* (Picossi et al., 2015) indicates that these regulatory changes likely contribute to the observed impairment.

In conclusion, deletion of the global transcription factor PacR in *Anabaena* triggers heterocyst formation even in NO_3_^-^ containing medium, likely due to impaired NO_3_^-^ uptake and disrupted NH_4_^+^ assimilation in the GOGAT-cycle (Fig. 5). This phenotype may be exacerbated by reduced PSI-yield and reduced expression of ferredoxin, which may lead to less reducing equivalents for nitrogen uptake. These results highlight the involvement of PacR in regulating nitrogen metabolism.

## Author Contributions

YA conceptualized the project. EW performed all experiments and wrote the first draft of the manuscript. TH, ASS, and LN supervised the experiments. All authors contributed to editing and proofreading. All authors have read and approved the final manuscript.

## Acknowledgements

We thank Tor Laurén at Åbo Akademi University for his assistance with elemental analysis, and Anni Nieminen, John Finell and Toveann Ahlnäs from the Institute for Molecular Medicine Finland (FIMM) for providing metabolic measurements. We thank Janne Isojärvi, Bradley Koch, Fiona Lynch, Peter Gollan and the Chipster-Team for their assistance with analyzing and graphing RNA sequencing data.

## Data Availability Statement

The RNA sequencing data used for Fig. 4,5 and Suppl. Tables 4-8 is available in NCBI with the GEO number GSE280386. Data supporting the findings of this study is available in the manuscript or supplementary data. Microscopic images and physiological source data is available on request.

