## Supporting Information for "The role of the LysR-type transcription factor PacR in regulating nitrogen metabolism in *Anabaena* sp. PCC7120"

Including:

Supplementary Figures 1-5

Supplementary Tables 1-8

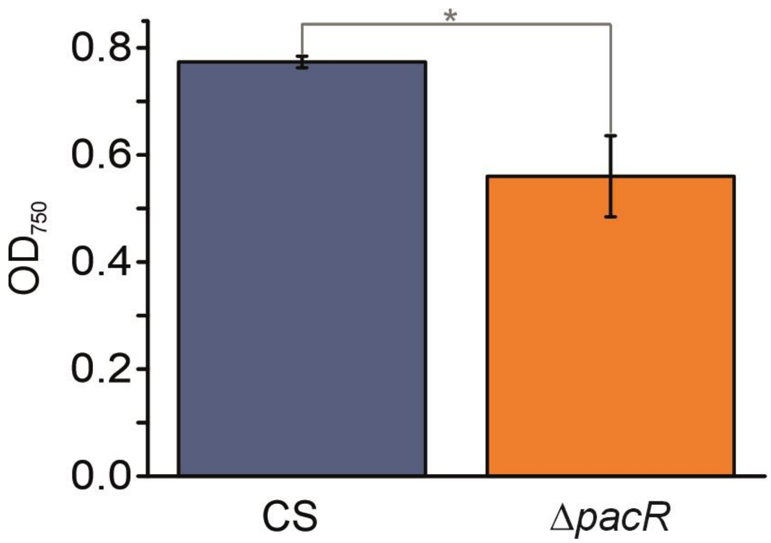

**Supplementary Figure 1.** Growth phenotype of the Δ*pacR* mutant in the presence of $NH_4^+$ monitored by $OD_{750}$ after 48 h in medium supplemented with 3 mM $NH_4^+$. Values are means ± SD; *n*=3 biologically independent experiments; statistical significance ($p<0.05$) is indicated by an asterisk.

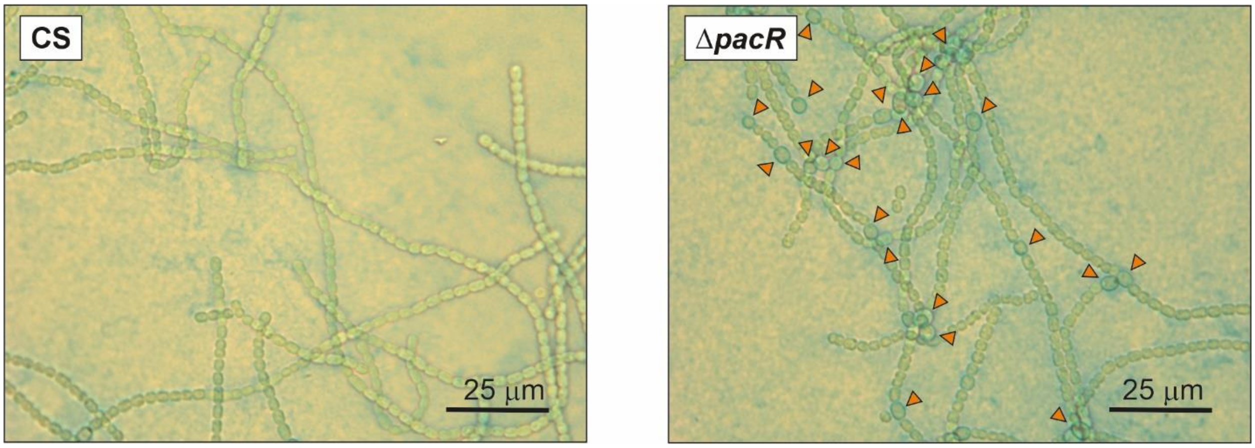

**Supplementary Figure 2.** Representative light-microscopy images of the CS and the Δ*pacR* mutant 48h after the shift from medium with $NH_4^+$ to medium with $NO_3^-$ as the sole nitrogen source. Arrows indicate heterocysts.

**Supplementary Table 1.** Heterocyst frequency for the CS and Δ*pacR* strains 48 h after the shift from medium with $NH_4^+$ to medium with $NH_4^+$ or $NO_3^-$ as the sole nitrogen source.

| **strain** | **nitrogen source of experimental culture** | **total # of cells counted** | **# of vegetative cells** | **# of heterocysts** | **heterocyst frequency [% of total cells counted]** |
|---|---|---|---|---|---|
| CS | $NH_4^+$ | 4379 | 4377 | 2 | 0.05±0.04 |
| | $NO_3^-$ | 5709 | 5655 | 54 | 0.95±0.16 |
| Δ*pacR* | $NH_4^+$ | 3754 | 3744 | 10 | 0.26±0.36[(1)] |
| | $NO_3^-$ | 5102 | 4722 | 380 | 7.45±1.68 |

[(1)] higher error due to occurrence of some heterocysts only in one biological replicate

**Supplementary Table 2.** N and C content of the CS and Δ*pacR* strains 48 h after the shift from medium with $NH_4^+$ to medium with $NO_3^-$ as the sole nitrogen source. Values are means ± SD; *n*=3 biologically independent experiments.

| strain | Nitrogen [weight %] | Carbon [weight %] |
|---|---|---|
| CS | 11.9±0.2 | 49.3±0.2 |
| Δ *pacR* | 10.2±0.8 | 47.6±0.8 |

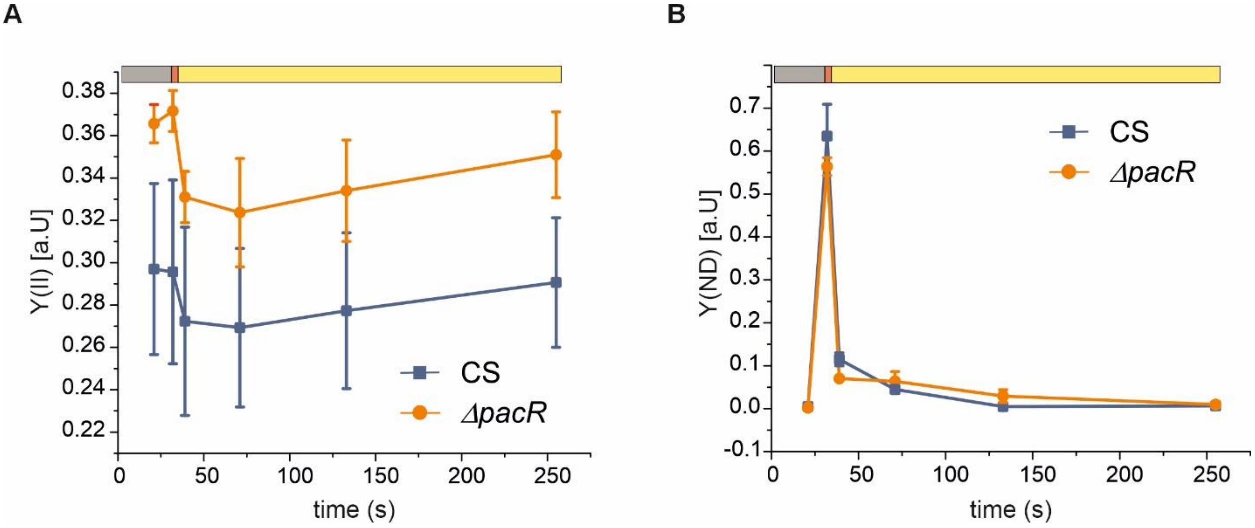

**Supplementary Figure 3.** Photosynthetic performance of the Δ*pacR* mutant and the CS 48 h after the shift from medium with $NH_4^+$ to medium with $NO_3^-$ as the sole nitrogen source. **A.** the effective yield of PSII [Y(II)] and **B.** the donor side limitation of PSI [Y(ND). black bar: darkness, red bar: far red light, yellow bar: actinic light. Values are mean ± SD, n=3 biological replicates.

**Supplementary Table 3.** The rates of $CO_2$ and $O_2$ exchange in the CS and Δ*pacR* mutant 48 h after the shift from $NH_4^+$ to $NO_3^-$ as sole nitrogen source. Gas exchange rates are presented as µmol mg Chl a-1 h-1. and values are mean ± SD, n =3 biological replicates. Asterisks indicate statistically significant differences compared to the CS (t-test, $p < 0.05$).

| | **Dark Respiration** | **Gross $O_2$ evolution (steady-state)** | **Light-induced $O_2$ uptake (max. peak / steady-state)** | **Net $O_2$ evolution (steady state)** | **$CO_2$ uptake (max. peak / steady-state)** |
|---|---|---|---|---|---|
| CS | 8.9±0.9 | 173.9±29.6 | 40.9±3.9* / 23.1±2.7* | 142.1±26.1 | 342.7±71.4 / 149.6±50.2 |
| Δ*pacR* | 13.5±3.7 | 179.3±12.5 | 13.0±3.0* / 7.3±5.0* | 159.4±11.1 | 424.6±36.7 / 173.5±34.9 |

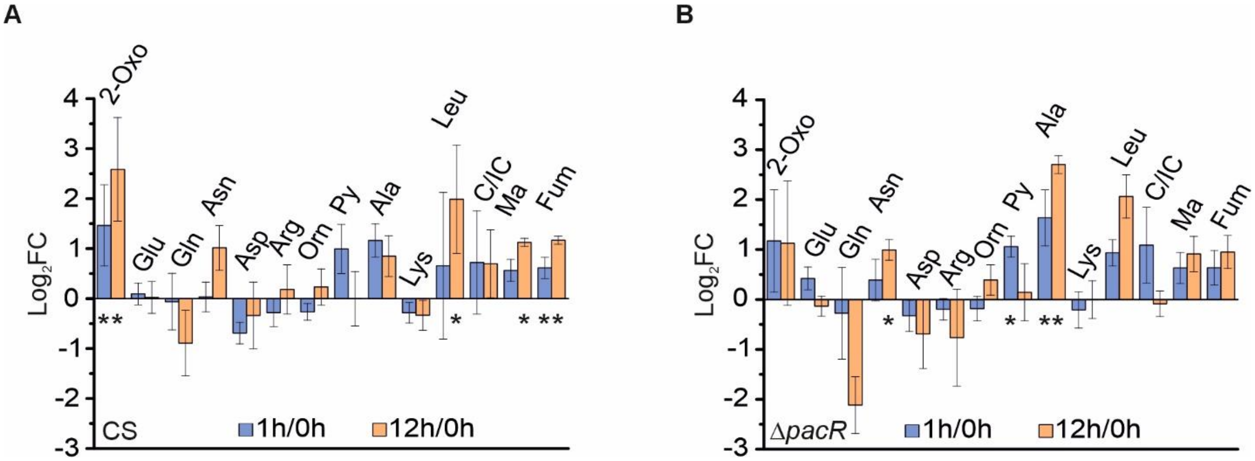

**Supplementary Figure 4.** Metabolite levels in the CS and Δ*pacR* mutant before and 1 h/12 h after the shift from $NH_4^+$ to $NO_3^-$ as sole nitrogen source. **A.** CS **B.** Δ*pacR*. Values of ion count/total ion count for t=1 h and t=12 h are in relation to t=0 h . 2-OG=2-Oxoglutarate, Glu=Glutamate, Gln=Glutamine, Asn=Asparagine, Asp=Aspartic acid, Arg=Arginine, Orn=Ornithine, Py=Pyruvate, Ala=Alanine, Lys=Lysine, Leu=Leucine, CI/C=Citrate/Isocitrate, Ma=Malic acid, Fum=Fumarate. Values are Mean ± SD, n=3-6 biological replicates shown as $log_2FC$. Asterisks indicate statistically significant differences, calculated based on differences in values of ion count/total ion count (t-test, $p < 0.05$).

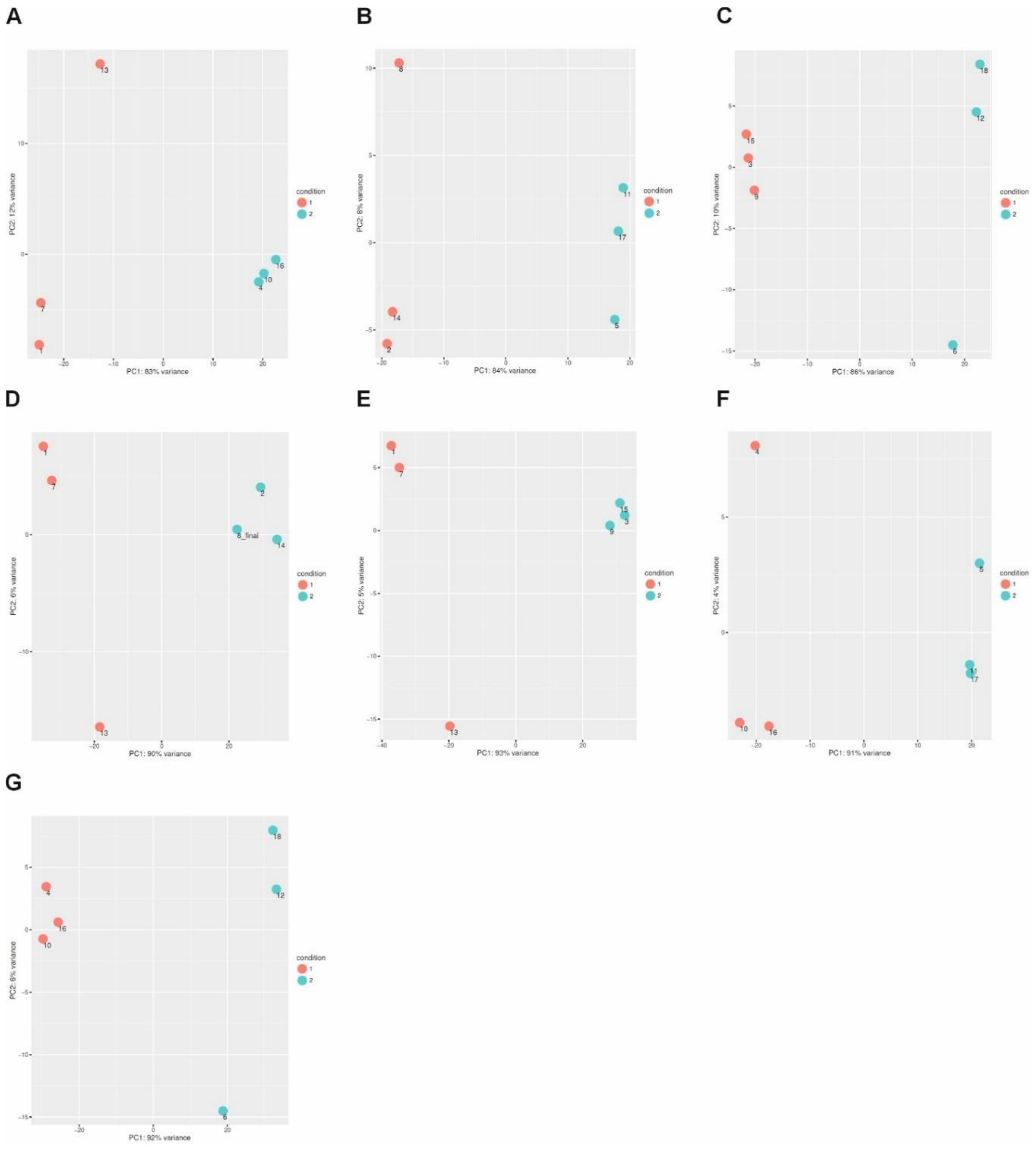

**Supplementary Figure 5.** PCA plots generated via the DESeq2 Bioconductor package for experimental level quality control of the RNA-seq data. **A.**, **B.** and **C.** show PCA plots for differential expression analysis for ∆*pacR* in relation to the CS. **D.** and **E.** show PCA plots for the differential expression analysis of the CS for t=1 h or 12 h in relation to 0 h. **F.** and **G.** show PCA plots for the differential expression analysis of ∆*pacR* for t=1 h or 12 h in relation to 0 h. The samples are numbered as follows: 1,7,13: 3 biological replicates for the CS at t=0 h; 4,10,16: 3 biological replicates for ∆*pacR* at t=0 h; 2,8,14: 3 biological replicates for the CS at t=1 h; 5,11,17: 3 biological replicates for ∆*pacR* at t=1 h; 3,9,15: 3 biological replicates for the CS at t=12 h; 6,12,18: 3 biological replicates for ∆*pacR* at t=12 h.

**Supplementary Table 4.** Differentially expressed genes at t=0, $log_2$FC ($\Delta$*pacR*/CS).

| **Category** | **ID** | **ENSEMBL_ID** | **Cyanobase _ID** | **$log_2$FC ($\Delta$pacR/CS)** |
|---|---|---|---|---|
| **N-metabolism** | | | | |
| Leucine dehydrogenase | *ldh* | *ENSB:DCJLnD1-vRm8aTd* | *all0426* | -1.45 |
| Cyanophycinase | *cphB_1* | *ENSB:qnAFH-zZi1_Cg7h* | *all0571* | -1.44 |
| Urease accessory protein | *ureG* | *ENSB:thD7sRcdm3nPz9H* | *alr0735* | 2.16 |
| Glutaminase 1 | *glsA1* | *ENSB:fjcIlBqee15rLqZ* | *all2934* | 2.08 |
| Molybdopterin-guanine dinucleotide biosynthesis protein A | *mobA* | *ENSB:P-Yyl0n7CNfjZCd* | *all0961* | 1.2 |
| Diaminobutyrate-pyruvate transaminase | *dat* | *ENSB:9sHAdEXZrAhlq7B* | *all0396* | -1.14 |
| Acetolactate synthase large subunit | *ilvB* | ENSB:MHH21VZ1XwX61y y | *all0427* | -1.25 |
| Proton/sodium-glutamate symport protein | *dctA* | ENSB:TZqUO6daj0fFv6c | *all0342* | -1.08 |
| Glycerate dehydrogenase | N/A | ENSB:zX_Ld1TTIPm3PX7 | *all8087* | -2.56 |

| | | | | |
|---|---|---|---|---|
| **Photosynthesis and respiration** | | | | |
| Homologue of the N-terminal domain of OCP | N/A | ENSB:Pjo4r6jSQqj9ZKS | *alr4783* | -1.22 |
| Homolog of the C-terminal domain of the OCP | N/A | ENSB:-Skx2hMko_CWcds | *all4940* | -1.17 |
| Bilin biosynthesis protein PecE | *pecE* | ENSB:QZBFZRurZXLurvC | *alr0526* | -1.52 |
| Bilin biosynthesis protein PecF | *pecF* | ENSB:zGBsC3D8Gtj2ymD | *alr0527* | -1.58 |
| Ferredoxin-1 | *petF* | ENSB:cn6HVgj9et9zpgG | *all4148* | -1.49 |
| Diflavin flavoprotein Flv4 | *flv4* | ENSB:dd_Ejl4qMz5TVdO | *all4446* | -1.48 |
| Pentapeptide repeat protein | N/A | ENSB:U7IgWWVcVHHMram | *alr5209* | -1.29 |
| Two-component response regulator | N/A | ENSB:wQizH64rRL3JSms | *alr0072* | -1.01 |
| **C-metabolism** | | | | |
| Sbta bicarbonate Na+ symporter | *sbtA* | ENSB:Z3aL43jCkgjvSJl | *all2134* | -1.83 |

| | | | | |
|---|---|---|---|---|
| Sucrose synthase | *susB* | ENSB:49m5v0JcFmdYuMU | *all1059* | -1.16 |
| Alpha,alpha-trehalase | *treH* | ENSB:lAnWugFkoPK758N | *all0166* | -1.43 |
| Malto-oligosyltrehalose trehalohydrolase | *treZ* | ENSB:wYIPo5aOxCDd1v_ | *all0168* | -1.65 |
| Transketolase | *tkt_2* | ENSB:bvaFhPk4hqCTUJt | *all4052* | -1.51 |
| Alpha-glucanotransferase | N/A | ENSB:8vPod86j-yWJPGD | *all0875* | -1.32 |
| **Phosphor-metabolism** | | | | |
| Phosphonate ABC transporter permease | N/A | ENSB:m4s0rAAffSgr4y7 | *all8088* | -4.75 |
| Phosphonate ABC transporter, phosphate-binding protein | N/A | ENSB:PbMDqwXP9BtwdTI | *all8089* | -5.41 |
| Phosphonate ABC transporter, ATP-binding component | *phnL* | ENSB:LSu_SJZWdPPP-RV | *all2217* | -1.3 |
| Phosphonate ABC transporter permease protein | *phnE* | ENSB:e6bTBbIW4RXbv2I | *all2227* | -1.32 |

| | | | | |
|---|---|---|---|---|
| ABC transporter, phosphate-binding protein; PhnD | *phnD* | ENSB:eQSxkgHwKX25u-K | *all2228* | -1.03 |
| ABC tranporter, ATP-binding component | *phnC1* | ENSB:bURWmyKBm4Qc7CU | *all2230* | -1.9 |
| Phosphodiesterase/alkaline phosphatase D | *phoD* | ENSB:hgDoJVzVX8lePKi | *alr2234* | -1.45 |
| **Metals** | | | | |
| Cation-efflux system membrane protein | N/A | ENSB:jOF_myIngWua0Ez | *all2845* | -1.37 |
| Similar to Na+/H+ antiporter | N/A | ENSB:9_uVQJPEWe0wqUI | *all4832* | -1.22 |
| ABC transporter, ATP-binding protein | *mntA* | ENSB:PmuYV0bqwlKOUDW | *all3575* | -1.07 |
| **Transcription factors** | | | | |
| PacR | *rbcR* | ENSB:3znthW3skDOmleL | *all3953* | -4.53 |

**Supplementary Table 5.** Differentially expressed genes at t=1, $log_2FC$ ($\Delta$*pacR*/CS).

| **Category** | **ID** | **ENSEMBL_ID** | **Cyanobase _ID** | **$log_2FC$ ($\Delta$pacR/CS)** |
|---|---|---|---|---|
| **N-metabolism** | | | | |
| Urease accessory protein | *ureG* | ENSB:thD7sRcdm3nPz9H | *alr0735* | 2.43 |
| Glutaminase 1 | *glsA1* | ENSB:fjcIlBqee15rLqZ | *all2934* | 1.54 |
| Molybdopterin-guanine dinucleotide biosynthesis protein A | *mobA* | ENSB:P-Yyl0n7CNfjZCd | *all0961* | 1.08 |
| Diaminobutyrate-pyruvate transaminase | *dat* | ENSB:9sHAdEXZrAhlq7B | *all0396* | -3.29 |
| Proton/sodium-glutamate symport protein | *dctA* | ENSB:TZqUO6daj0fFv6c | *all0342* | -2.24 |
| Nitrite reductase | *nirA* | ENSB:e44ECOSAHPRWrlu | *alr0607* | -1.52 |
| Nitrate transport nitrate-binding protein | *nrtA* | ENSB:V1D0skKAkHJgWSZ | *alr0608* | -1.44 |
| Nitrate transport permease protein | *nrtB* | ENSB:7v-sc3mUDbuJK5G | *alr0609* | -1.62 |
| Nitrate transport ATP-binding protein | *nrtC* | ENSB:Ys8OE0aTWeoXMXH | *alr0610* | -1.61 |

| | | | | |
|---|---|---|---|---|
| Nitrate transport ATP-binding protein | *nrtD* | ENSB:YD5wpg9hsgLEiVJ | *alr0611* | -1.62 |
| Nitrate reductase | *narB* | ENSB:EYOY4XLgkxMnQT5 | *alr0612* | -1.5 |
| **Photosynthesis and respiration** | | | | |
| NTD-OCP like protein | N/A | ENSB:YZqer7AeouGwj-a | *all4941* | -1.27 |
| Photosystem II protein D1 | *psbA3* | ENSB:9agEV78l86gFA_V | *alr4592* | -2.09 |
| Photosystem II protein D1 | *psbA1* | ENSB:yJKfo7_EpcR1xJq | *alr4866* | 1.68 |
| Light-independent protochlorophyllide reductase iron-sulfur ATP-binding protein | *chlL* | ENSB:O1DJvDa30jvN8hW | *all5078* | 1.06 |
| Ferredoxin-1 | *petF* | ENSB:cn6HVgj9et9zpgG | *all4148* | -1.99 |
| Diflavin flavoprotein Flv2 | *flv2* | ENSB:txsyY6CJmI0pCJY | *all4444* | -1.29 |
| Diflavin flavoprotein Flv4 | *flv4* | ENSB:dd_Ejl4qMz5TVdO | *all4446* | -3.34 |
| Pentapeptide repeat protein | N/A | ENSB:U7IgWWVcVHHMram | *alr5209* | -1.43 |

| | | | | |
|---|---|---|---|---|
| **C-metabolism** | | | | |
| Sbta bicarbonate Na+ symporter | *sbtA* | ENSB:Z3aL43jCkgjvSJl | *all2134* | -4.33 |
| Bicarbonate transport system permease protein | *cmpB* | ENSB:Q5MPpHiTaAnt52i | *alr2878* | -1.28 |
| **Transcription Factors** | | | | |
| PacR | *rbcR* | ENSB:3znthW3skDOmleL | *all3953* | -4.62 |
| Cell wall-binding protein Fur | *furA* | ENSB:D3Qm5IOjvV5LGG N | *all1691* | 1.14 |

**Supplementary Table 6.** Differentially expressed genes at t=12, $log_2$FC (Δ*pacR*/CS)

| **Category** | **ID** | **ENSEMBL_ID** | **Cyanobase _ID** | **$log_2$FC (Δ*pacR*/CS)** |
|---|---|---|---|---|
| **N-metabolism** | | | | |
| Urease accessory protein | *ureG* | *ENSB:thD7sRcdm3nPz9H* | *alr0735* | 2.17 |
| Glutaminase 1 | *glsA1* | *ENSB:fjcIlBqee15rLqZ* | *all2934* | 1.51 |
| Molybdopterin-guanine dinucleotide biosynthesis protein A | *mobA* | *ENSB:P-Yyl0n7CNfjZCd* | *all0961* | 1.11 |
| Diaminobutyrate-pyruvate transaminase | *dat* | *ENSB:9sHAdEXZrAhlq7B* | *all0396* | -1.03 |
| Alanine dehydrogenase | *ald* | *ENSB:rWYGwN8UWkGzo2q* | *alr2355* | 1.24 |
| Nitrogen stress-induced RNA 1 | N/A | *ENSB:7BhRYdNgWHwPQX0* | N/A | 2.08 |
| Nitrogen stress-induced RNA 1 | N/A | *ENSB:kLcNZRN7-pJLBxt* | N/A | 1.7 |
| Nitrogen stress-induced RNA 1 | N/A | *ENSB:YhC5bG0zn0YQBkL* | N/A | 1.63 |
| **Heterocyst formation** | | | | |
| Glycolipid synthase | *hglT* | *ENSB:5ThqEjI9sShuE-E* | *all5341* | 3.27 |

| | | | | |
|---|---|---|---|---|
| Ketoacyl reductase | *hetN* | *ENSB:ZMQxbwAIyd9Q-fM* | *alr5358* | 1.28 |
| Glucosyltransferase | N/A | *ENSB:RmnPuja33XjbEKh* | *all2289* | 1 |
| Heterocyst envelope polysaccharide synthesis protein | *hepD* | *ENSB:Oc1V-X3m6qCq5eW* | *alr3698* | 3.85 |
| Glycosyltransferase | *hepE* | *ENSB:94sxNIFAyRkqwlV* | *alr3699* | 1.43 |
| Glucose-1-phosphate cytidylyltransferase | N/A | *ENSB:ZjfrPap_6rDqJvJ* | *alr2825* | 4.91 |
| Glycosyltransferase | N/A | *ENSB:aofJPhZKhvBy41C* | *alr2836* | 3.39 |
| Glycosyltransferase | N/A | *ENSB:ffiqyvkzMYyafmg* | *alr2839* | 3.38 |
| dTDP-4-dehydrorhamnose 3,5-epimerase | *rfbC* | *ENSB:CNlG9yMspfYs3EQ* | *alr2830* | 4.09 |
| Heterocyst differentiation ATP-binding protein | *hepA* | *ENSB:4TIFljsR-svhXZJ* | *alr2835* | 3.49 |
| Nitrogen fixation protein | *nifW* | *ENSB:KLm42LryWdoGB7J* | *all1433* | 5.33 |
| Nitrogenase molybdenum-iron protein beta chain | *nifN* | *ENSB:GgyBKW9hY4nEOAP* | *all1437* | 3.74 |
| Nitrogenase molybdenum-iron protein beta chain | *nifK* | *ENSB:p8L0nBpHJJRKNhu* | *all1440* | 6.61 |

| | | | | |
|---|---|---|---|---|
| Nitrogenase molybdenum-iron protein alpha chain | *nifD* | *ENSB:SBhFQ-pgKfU865y* | *all1454* | 4.78 |
| Nitrogenase iron protein | *nifH1* | *ENSB:-UiwXBMw740um_c* | *all1455* | 3.16 |
| Nitrogen fixation protein | *nifU* | *ENSB:zD7Me4dVDd5AIk-* | *all1456* | 2.9 |
| Nitrogenase cofactor synthesis protein | *nifS* | *ENSB:cwZOUWtsHDDQELT* | *all1457* | 3.64 |
| Homocitrate synthase | *nifV1* | *ENSB:xnTLVRDJMQvACq9* | *alr1407* | 3.65 |
| Nitrogen fixation protein | *nifB* | *ENSB:oKdYkJwJmedayE7* | *all1517* | 3.13 |
| Protein HesB, heterocyst | *hesB* | *ENSB:0W1BDOVJav4vemq* | *all1431* | 3.58 |
| Protein HesA, heterocyst | *hesA* | *ENSB:Byut0LlNHsEQVLj* | *all1432* | 2.65 |
| Periplasmic [NiFeSe] hydrogenase large subunit | *hupL* | *ENSB:q7HshiWY1tEwz49* | *all0687* | 3.5 |
| Periplasmic [NiFe] hydrogenase small subunit | *hupS* | *ENSB:YybVoSB-3p1afrt* | *all0688* | 2.93 |
| Ferredoxin | *fdxB* | *ENSB:g-hu8fN-yJ9QOkL* | *asr2513* | 2.56 |
| Ferredoxin, heterocyst | *fdxH* | *ENSB:pgoFbyUO25Y11KZ* | *all1430* | 3.18 |

| | | | | |
|---|---|---|---|---|
| Diflavin flavoprotein flv1B | *flv1B* | *ENSB:egKkM6AIKDL3lV3* | *all0177* | 1.3 |
| Putative diflavin flavoprotein flv3B | *flv3B* | *ENSB:gyAx2_YtlGWpP0r* | *all0178* | 1.05 |
| **C-metabolism** | | | | |
| Sbta bicarbonate Na+ symporter | *sbtA* | *ENSB:Z3aL43jCkgjvSJl* | *all2134* | -1.08 |
| 6-Phosphofructokinase | *pfkA1* | *ENSB:STpcob1Kje9jW0R* | *all7335* | -1.11 |
| Galactosyltransferase | N/A | *ENSB:F5rC4qxkMvFfiwq* | *all2037* | 1.78 |
| **Photosynthesis and respiration** | | | | |
| Phycoerythrocyanin subunit beta | *pecB* | *ENSB:DlJg8tfpJb3Pc02* | *alr0523* | -1.98 |
| Phycoerythrocyanin alpha chain | *pecA* | *ENSB:DkyiMvYmj7wtDsq* | *alr0524* | -1.85 |
| Phycobilisome 34.5 kDa linker polypeptide, phycoerythrocyanin-associated, rod | *pecC* | *ENSB:4P_EwWN8WSSWyss* | *alr0525* | -2.06 |
| Bilin biosynthesis protein PecE | *pecE* | *ENSB:QZBFZRurZXLurvC* | *alr0526* | -2.68 |

| | | | | |
|---|---|---|---|---|
| Bilin biosynthesis protein PecF | *pecF* | *ENSB:zGBsC3D8Gtj2ymD* | *alr0527* | -2.46 |
| Photosystem II CP43 reaction center protein homologue | *isiA2* | *ENSB:n2bv19CnTlJ6ejC* | *all4002* | 1.59 |
| Photosystem II CP43 reaction center protein homologue | *isiA3* | *ENSB:72J4WLJgrkah1ix* | *all4003* | 2.06 |
| Oxygen-independent coproporphyrinogen III oxidase | *hemN* | *ENSB:Klj8zn66_ssKF-m* | *alr3126* | 1.03 |
| Ferredoxin-1 | *petF* | *ENSB:cn6HVgj9et9zpgG* | *all4148* | -2.23 |
| Flavodoxin I | *isiB* | *ENSB:vCA6KyCuxG-PG7-* | *alr2405* | 1.94 |
| Cytochrome c oxidase subunit 1 | *coxA2* | *ENSB:6VxYhoGYWmvmjDB* | *alr2515* | 3.22 |
| Putative cytochrome c oxidase subunit 3 | *coxC2* | *ENSB:Oyzg-naQZMZT0Ld* | *alr2516* | 3.67 |
| Cytochrome c oxidase subunit 1 | *coxA3* | *ENSB:OjIeYMGIKj96usn* | *alr2732* | 2.44 |
| Oxidoreductase | N/A | *ENSB:_N-AKaQFkqo5q49* | *all5345* | 3.28 |
| Pentapeptide repeat protein | N/A | *ENSB:U7IgWWVcVHHMram* | *alr5209* | -1.43 |
| **Phosphor-metabolism** | | | | |

| | | | | |
|---|---|---|---|---|
| Phosphonate ABC transporter, ATP-binding component | *phnK* | *ENSB:L4cUHY-SreLDCy2* | *all2218* | -1.01 |
| ABC transporter, periplasmic phosphate-binding protein | N/A | *ENSB:ANyliMgnrcBMIfb* | *alr1094* | 2.18 |
| **Transcription factors** | | | | |
| PacR | *rbcR* | *ENSB:3znthW3skDOmleL* | *all3953* | -4.85 |
| Nitrogen-responsive regulatory protein | *ntcA* | *ENSB:wHiGICXo9Og7Xlz* | *alr4392* | 1.12 |
| RNA polymerase sigma-subunit | *sigC* | *ENSB:WGvPMBCl6Hl3Gwg* | *all1692* | 1.1 |
| AraC type Transcriptional regulator | N/A | *ENSB:f_q6Xin8pCUNvPv* | *all2035* | 1.03 |

**Supplementary Table 7.** Differentially expressed genes for the CS, $log_2FC$ (t1/t0) and $log_2FC$ (t12/t0)

| **Category** | **ID** | **ENSEMBL_ID** | **Cyanobase _ID** | **$log_2FC$ (t1/t0)** | **$log_2FC$ (t12/t0)** |
|---|---|---|---|---|---|
| **N-metabolism** | | | | | |
| Leucine dehydrogenase | *ldh* | *ENSB:DCJLnD1-vRm8aTd* | *all0426* | -1.01 | -2.45 |
| Cyanophycinase | *cphB _1* | *ENSB:qnAFH-zZi1_Cg7h* | *all0571* | -4.01 | -4.51 |
| Urease accessory protein | *ureG* | *ENSB:thD7sRcdm3nPz9H* | *alr0735* | N/A | N/A |
| Glutaminase 1 | *glsA1* | *ENSB:fjcIlBqee15rLqZ* | *all2934* | N/A | N/A |
| Molybdopterin-guanine dinucleotide biosynthesis protein A | *mobA* | *ENSB:P-Yyl0n7CNfjZCd* | *all0961* | N/A | N/A |
| Diaminobutyrate-pyruvate transaminase | *dat* | *ENSB:9sHAdEXZrAhlq7B* | *all0396* | N/A | N/A |
| Acetolactate synthase large subunit | *ilvB* | *ENSB:MHH21VZ1XwX61y y* | *all0427* | -1.08 | -1.86 |
| Proton/sodium-glutamate symport protein | *dctA* | *ENSB:TZqUO6daj0fFv6c* | *all0342* | N/A | N/A |
| Glycerate dehydrogenase | N/A | *ENSB:zX_Ld1TTIPm3PX7* | *all8087* | -3.45 | -2.36 |

| | | | | | |
|---|---|---|---|---|---|
| Alanine dehydrogenase | *ald* | *ENSB:rWYGwN8UWkGzo2q* | *alr2355* | N/A | N/A |
| Glutamine synthetase | *glnA* | *ENSB:ab0fhVsYEpHjH88* | *alr2328* | N/A | 2.11 |
| Nitrate transport nitrate-binding protein | *nrtA* | *ENSB:V1D0skKAkHJgWSZ* | *alr0608* | 4.58 | 5.37 |
| Nitrate transport permease protein | *nrtB* | *ENSB:7v-sc3mUDbuJK5G* | *alr0609* | 4.55 | 5.15 |
| Nitrate transport ATP-binding protein | *nrtC* | *ENSB:Ys8OE0aTWeoXMXH* | *alr0610* | 4.49 | 4.94 |
| Nitrate transport ATP-binding protein | *nrtD* | *ENSB:YD5wpg9hsgLEiVJ* | *alr0611* | 4.45 | 4.88 |
| Nitrate reductase | *narB* | *ENSB:EYOY4XLgkxMnQT5* | *alr0612* | 3.04 | 3.44 |
| Nitrite reductase | *nirA* | *ENSB:e44ECOSAHPRWrlu* | *alr0607* | 4.59 | 5.43 |
| Nitrogen stress-induced RNA 1 | N/A | *ENSB:7BhRYdNgWHwPQX0* | N/A | N/A | 1.74 |
| Nitrogen stress-induced RNA 1 | N/A | *ENSB:kLcNZRN7-pJLBxt* | N/A | N/A | N/A |
| Nitrogen stress-induced RNA 1 | N/A | *ENSB:YhC5bG0zn0YQBkL* | N/A | N/A | 1.04 |
| **Heterocyst formation** | | | | | |

| | | | | | |
|---|---|---|---|---|---|
| Glycolipid synthase | *hglT* | *ENSB:5ThqEjI9sShuE-E* | *all5341* | N/A | N/A |
| Ketoacyl reductase | *hetN* | *ENSB:ZMQxbwAIyd9Q-fM* | *alr5358* | N/A | N/A |
| Glucosyltransferase | N/A | *ENSB:RmnPuja33XjbEKh* | *all2289* | N/A | 1.96 |
| Heterocyst envelope polysaccharide synthesis protein | *hepD* | *ENSB:Oc1V-X3m6qCq5eW* | *alr3698* | N/A | N/A |
| Glycosyltransferase | *hepE* | *ENSB:94sxNIFAyRkqwlV* | *alr3699* | N/A | N/A |
| Glucose-1-phosphate cytidylyltransferase | N/A | *ENSB:ZjfrPap_6rDqJvJ* | *alr2825* | N/A | 1.3 |
| Glycosyltransferase | N/A | *ENSB:aofJPhZKhvBy41C* | *alr2836* | N/A | N/A |
| Glycosyltransferase | N/A | *ENSB:ffiqyvkzMYyafmg* | *alr2839* | N/A | 1.03 |
| dTDP-4-dehydrorhamnose 3,5-epimerase | *rfbC* | *ENSB:CNlG9yMspfYs3EQ* | *alr2830* | N/A | 1.07 |
| Heterocyst differentiation ATP-binding protein | *hepA* | *ENSB:4TIFljsR-svhXZJ* | *alr2835* | N/A | N/A |
| Nitrogen fixation protein | *nifW* | *ENSB:KLm42LryWdoGB7J* | *all1433* | N/A | N/A |
| Nitrogenase molybdenum iron protein beta chain | *nifN* | *ENSB:GgyBKW9hY4nEOAP* | *all1437* | N/A | N/A |

| | | | | | |
|---|---|---|---|---|---|
| Nitrogenase molybdenum iron protein beta chain | *nifK* | *ENSB:p8L0nBpHJJRKNhu* | *all1440* | N/A | N/A |
| Nitrogenase molybdenum iron protein alpha chain | *nifD* | *ENSB:SBhFQ-pgKfU865y* | *all1454* | N/A | N/A |
| Nitrogenase iron protein | *nifH1* | *ENSB:-UiwXBMw740um_c* | *all1455* | N/A | N/A |
| Nitrogen fixation protein | *nifU* | *ENSB:zD7Me4dVDd5AIk-* | *all1456* | N/A | N/A |
| Nitrogenase cofactor synthesis protein | *nifS* | *ENSB:cwZOUWtsHDDQELT* | *all1457* | N/A | N/A |
| Homocitrate synthase | *nifV1* | *ENSB:xnTLVRDJMQvACq9* | *alr1407* | N/A | N/A |
| Nitrogen fixation protein | *nifB* | *ENSB:oKdYkJwJmedayE7* | *all1517* | N/A | N/A |
| Protein HesB, heterocyst | *hesB* | *ENSB:0W1BDOVJav4vemq* | *all1431* | N/A | N/A |
| Protein HesA, heterocyst | *hesA* | *ENSB:Byut0LlNHsEQVLj* | *all1432* | 1.83 | 2.92 |
| Periplasmic [NiFeSe] hydrogenase large subunit | *hupL* | *ENSB:q7HshiWY1tEwz49* | *all0687* | N/A | N/A |
| Periplasmic [NiFe] hydrogenase small subunit | *hupS* | *ENSB:YybVoSB-3p1afrt* | *all0688* | N/A | N/A |

| | | | | | |
|---|---|---|---|---|---|
| Ferredoxin | *fdxB* | *ENSB:g-hu8fN-yJ9QOkL* | *asr2513* | N/A | N/A |
| Ferredoxin, heterocyst | *fdxH* | *ENSB:pgoFbyUO25Y11KZ* | *all1430* | N/A | N/A |
| Diflavin flavoprotein flv1B | *flv1B* | *ENSB:egKkM6AIKDL3lV3* | *all0177* | N/A | N/A |
| Diflavin flavoprotein flv3B | *flv3B* | *ENSB:gyAx2_YtlGWpP0r* | *all0178* | N/A | N/A |
| **Photosynthesis and respiration** | | | | | |
| NTD-OCP like protein | N/A | *ENSB:YZqer7AeouGwj-a* | *all4941* | 1.89 | -1.62 |
| Homologue of the N-terminal domain of OCP | N/A | *ENSB:Pjo4r6jSQqj9ZKS* | *alr4783* | -3.58 | -3.53 |
| Homolog of the C-terminal domain of the OCP | N/A | *ENSB:-Skx2hMko_CWcds* | *All4940* | -2.18 | -2.11 |
| Phycoerythrocyanin subunit beta | *pecB* | *ENSB:DlJg8tfpJb3Pc02* | *alr0523* | -3.9 | N/A |
| Phycoerythrocyanin alpha chain | *pecA* | *ENSB:DkyiMvYmj7wtDsq* | *alr0524* | -3.97 | N/A |
| Phycobilisome 34.5 kDa linker polypeptide, phycoerythrocyanin-associated, rod | *pecC* | *ENSB:4P_EwWN8WSSWyss* | *alr0525* | -4.41 | N/A |

| | | | | | |
|---|---|---|---|---|---|
| Bilin biosynthesis protein | *pecE* | *ENSB:QZBFZRurZXLurvC* | *alr0526* | -4.65 | N/A |
| Bilin biosynthesis protein | *pecF* | *ENSB:zGBsC3D8Gtj2ymD* | *alr0527* | -3.69 | N/A |
| Photosystem II protein D1 | *psbA 3* | *ENSB:9agEV78l86gFA_V* | *alr4592* | 2.61 | -1.82 |
| Photosystem II protein D1 | *psbA 1* | *ENSB:yJKfo7_EpcR1xJq* | *alr4866* | N/A | N/A |
| Photosystem II CP43 reaction center protein homologue | *isiA2* | *ENSB:n2bv19CnTlJ6ejC* | *all4002* | N/A | N/A |
| Photosystem II CP43 reaction center protein homologue | *isiA3* | *ENSB:72J4WLJgrkah1ix* | *all4003* | N/A | N/A |
| Light-independent protochlorophyllide reductase iron-sulfur ATP-binding protein | *chlL* | *ENSB:O1DJvDa30jvN8hW* | *all5078* | -2.65 | N/A |
| Oxygen-independent coproporphyrinogen III oxidase | *hemN* | *ENSB:Klj8zn66_ssKF-m* | *alr3126* | N/A | N/A |
| Ferredoxin-1 | *petF* | *ENSB:cn6HVgj9et9zpgG* | *all4148* | N/A | N/A |
| Flavodoxin | *isiB* | *ENSB:vCA6KyCuxG-PG7-* | *alr2405* | N/A | N/A |
| Diflavin flavoprotein flv2 | *flv2* | *ENSB:txsyY6CJmI0pCJY* | *all4444* | 1.01 | N/A |

| | | | | | |
|---|---|---|---|---|---|
| Diflavin flavoprotein Flv4 | *flv4* | *ENSB:dd_Ejl4qMz5TVdO* | *all4446* | N/A | -1.14 |
| Cytochrome c oxidase subunit 1 | *CoxA2* | *ENSB:6VxYhoGYWmvmjDB* | *alr2515* | N/A | N/A |
| Putative cytochrome c oxidase subunit 3 | *coxC2* | *ENSB:Oyzg-naQZMZT0Ld* | *alr2516* | N/A | N/A |
| Cytochrome c oxidase subunit 1 | *coxA3* | *ENSB:OjIeYMGIKj96usn* | *alr2732* | N/A | N/A |
| Oxidoreductase | N/A | *ENSB:_N-AKaQFkqo5q49* | *all5345* | N/A | N/A |
| Pentapeptide repeat protein | N/A | *ENSB:U7IgWWVcVHHMram* | *alr5209* | N/A | N/A |
| Two-component response regulator | N/A | *ENSB:wQizH64rRL3JSms* | *alr0072* | -2.1 | -1.99 |
| **C-metabolism** | | | | | |
| Sbta bicarbonate Na+ symporter | *sbtA* | *ENSB:Z3aL43jCkgjvSJl* | *all2134* | 2.27 | N/A |
| Bicarbonate transport system permease protein | *cmpB* | *ENSB:Q5MPpHiTaAnt52i* | *alr2878* | 1.19 | N/A |
| Sucrose synthase | *susB* | *ENSB:49m5v0JcFmdYuMU* | *all1059* | -4.13 | -4.2 |
| Alpha,alpha- trehalase | *treH* | *ENSB:lAnWugFkoPK758N* | *all0166* | -3.55 | -4.02 |

| | | | | | |
|---|---|---|---|---|---|
| Malto-oligosyltrehalose trehalohydrolase | *treZ* | *ENSB:wYIPo5aOxCDd1v_* | *all0168* | -4.28 | -5.15 |
| Transketolase | *tkt_2* | *ENSB:bvaFhPk4hqCTUJt* | *all4052* | -4.3 | -4.68 |
| Alpha-glucanotransferase | N/A | *ENSB:8vPod86j-yWJPGD* | *all0875* | -5.6 | -6.42 |
| 6-Phosphofructokinase | *pfkA1* | *ENSB:STpcob1Kje9jW0R* | *all7335* | N/A | 1.23 |
| Galactosyltransferase | N/A | *ENSB:F5rC4qxkMvFfiwq* | *all2037* | N/A | N/A |
| **Metals** | | | | | |
| Cation-efflux system membrane protein | N/A | *ENSB:jOF_mylngWua0Ez* | *all2845* | -2.02 | -2.47 |
| Similar to Na+/H+ antiporter | N/A | *ENSB:9_uVQJPEWe0wqUI* | *all4832* | -2.59 | -3.01 |
| ABC transporter, ATP-binding protein | *mntA* | *ENSB:PmuYV0bqwlKOUDW* | *all3575* | N/A | -1.78 |
| **Phosphor-metabolism** | | | | | |
| Phosphonate ABC transporter permease | N/A | *ENSB:m4s0rAAffSgr4y7* | *all8088* | -4.21 | -4.33 |
| Phosphonate ABC transporter, | N/A | *ENSB:PbMDqwXP9BtwdTI* | *all8089* | -5.68 | -1.88 |

| | | | | | |
|---|---|---|---|---|---|
| phosphate-binding protein | | | | | |
| ATP-binding component | *phnL* | *ENSB:LSu_SJZWdPPP-RV* | *all2217* | -1.32 | -1.34 |
| phosphonate ABC transporter, ATP-binding component | *phnK* | *ENSB:L4cUHY-SreLDCy2* | *all2218* | N/A | N/A |
| phosphonate ABC transporter permease protein | *phnE* | *ENSB:e6bTBbIW4RXbv2I* | *all2227* | N/A | N/A |
| ABC tranporter, phosphate-binding protein; PhnD | *phnD* | *ENSB:eQSxkgHwKX25u-K* | *all2228* | -1.56 | -1.88 |
| ABC transporter, ATP-binding component | *phnC1* | *ENSB:bURWmyKBm4Qc7CU* | *all2230* | -3.93 | -3.4 |
| Phosphodiesterase/alkaline phosphatase D | *phoD* | *ENSB:hgDoJVzVX8lePKi* | *alr2234* | -2 | -2.89 |
| ABC transporter, periplasmic phosphate-binding protein | N/A | *ENSB:ANyliMgnrcBMIfb* | *alr1094* | N/A | N/A |
| **Transcription Factors** | | | | | |
| PacR | *rbcR* | *ENSB:3znthW3skDOmleL* | *all3953* | N/A | N/A |
| Nitrogen-responsive regulatory protein | *ntcA* | *ENSB:wHiGICXo9Og7Xlz* | *alr4392* | N/A | N/A |

| | | | | | |
|---|---|---|---|---|---|
| Cell wall-binding protein Fur | *furA* | *ENSB:D3Qm5IOjvV5LGGN* | *all1691* | -1.3 | N/A |
| RNA polymerase sigma-subunit | *sigC* | *ENSB:WGvPMBCl6Hl3Gwg* | *all1692* | N/A | 1.8 |
| AraC type Transcriptional regulator | N/A | *ENSB:f_q6Xin8pCUNvPv* | *all2035* | N/A | 3.13 |

**Supplementary Table 8.** Differentially expressed genes for Δ*pacR*, $log_2$FC (t1/t0) and $log_2$FC (t12/t0)

| Category | ID | ENSEMBL_ID | Cyanobase _ID | $log_2$FC (t1/t0) | $log_2$FC (t12/t0) |
|---|---|---|---|---|---|
| **N-metabolism** | | | | | |
| Leucine dehydrogenase | *ldh* | *ENSB:DCJLnD1-vRm8aTd* | *all0426* | N/A | N/A |
| Cyanophycinase | *cphB _1* | *ENSB:qnAFH-zZi1_Cg7h* | *all0571* | -2.36 | -2.96 |
| Urease accessory protein | *ureG* | *ENSB:thD7sRcdm3nPz9H* | *alr0735* | N/A | N/A |
| Glutaminase 1 | *glsA1* | *ENSB:fjcIlBqee15rLqZ* | *all2934* | N/A | N/A |
| Molybdopterin-guanine dinucleotide biosynthesis protein A | *mobA* | *ENSB:P-Yyl0n7CNfjZCd* | *all0961* | N/A | N/A |
| Diaminobutyrate-pyruvate transaminase | *dat* | *ENSB:9sHAdEXZrAhlq7B* | *all0396* | N/A | N/A |
| Acetolactate synthase large subunit | *ilvB* | *ENSB:MHH21VZ1XwX61yy* | *all0427* | N/A | N/A |
| Proton/sodium-glutamate symport protein | *dctA* | *ENSB:TZqUO6daj0fFv6c* | *all0342* | N/A | N/A |
| Glycerate dehydrogenase | N/A | *ENSB:zX_Ld1TTIPm3PX7* | *all8087* | N/A | N/A |

| | | | | | |
|---|---|---|---|---|---|
| Alanine dehydrogenase | *ald* | *ENSB:rWYGwN8UWkGzo2q* | *alr2355* | N/A | 1.52 |
| Glutamine synthetase | *glnA* | *ENSB:ab0fhVsYEpHjH88* | *alr2328* | 1 | 1.96 |
| Nitrate transport nitrate-binding protein | *nrtA* | *ENSB:V1D0skKAkHJgWSZ* | *alr0608* | 4.08 | 6.26 |
| Nitrate transport permease protein | *nrtB* | *ENSB:7v-sc3mUDbuJK5G* | *alr0609* | 3.84 | 5.98 |
| Nitrate transport ATP-binding protein | *nrtC* | *ENSB:Ys8OE0aTWeoXMXH* | *alr0610* | 3.97 | 5.93 |
| Nitrate transport ATP-binding protein | *nrtD* | *ENSB:YD5wpg9hsgLEiVJ* | *alr0611* | 4.06 | 6.01 |
| Nitrate reductase | *narB* | *ENSB:EYOY4XLgkxMnQT5* | *alr0612* | 2.09 | 3.92 |
| Nitrite reductase | *nirA* | *ENSB:e44ECOSAHPRWrlu* | *alr0607* | 4.07 | 6.49 |
| Nitrogen stress-induced RNA 1 | N/A | *ENSB:7BhRYdNgWHwPQX0* | N/A | N/A | 3.97 |
| Nitrogen stress-induced RNA 1 | N/A | *ENSB:kLcNZRN7-pJLBxt* | N/A | N/A | 1.86 |
| Nitrogen stress-induced RNA 1 | N/A | *ENSB:YhC5bG0zn0YQBkL* | N/A | N/A | 2.63 |
| **Heterocyst formation** | | | | | |

| | | | | | |
|---|---|---|---|---|---|
| Glycolipid synthase | *hglT* | *ENSB:5ThqEjI9sShuE-E* | *all5341* | N/A | 2.98 |
| Ketoacyl reductase | *hetN* | *ENSB:ZMQxbwAIyd9Q-fM* | *alr5358* | N/A | N/A |
| Glucosyltransferase | N/A | *ENSB:RmnPuja33XjbEKh* | *all2289* | N/A | 2.41 |
| Heterocyst envelope polysaccharide synthesis protein | *hepD* | *ENSB:Oc1V-X3m6qCq5eW* | *alr3698* | N/A | 3.89 |
| Glycosyltransferase | *hepE* | *ENSB:94sxNIFAyRkqwlV* | *alr3699* | N/A | 1.38 |
| Glucose-1-phosphate cytidylyltransferase | N/A | *ENSB:ZjfrPap_6rDqJvJ* | *alr2825* | N/A | 6.95 |
| Glycosyltransferase | N/A | *ENSB:aofJPhZKhvBy41C* | *alr2836* | N/A | 3.66 |
| Glycosyltransferase | N/A | *ENSB:ffiqyvkzMYyafmg* | *alr2839* | N/A | 4.3 |
| dTDP-4-dehydrorhamnose 3,5-epimerase | *rfbC* | *ENSB:CNlG9yMspfYs3EQ* | *alr2830* | N/A | 5.09 |
| Heterocyst differentiation ATP-binding protein HepA | *hepA* | *ENSB:4TIFljsR-svhXZJ* | *alr2835* | N/A | 4.08 |
| Nitrogen fixation protein | *nifW* | *ENSB:KLm42LryWdoGB7J* | *all1433* | N/A | 3.07 |
| Nitrogenase molybdenum-iron protein beta chain | *nifN* | *ENSB:GgyBKW9hY4nEOAP* | *all1437* | N/A | 3.87 |

| | | | | | |
|---|---|---|---|---|---|
| Nitrogenase molybdenum-iron protein beta chain | *nifK* | *ENSB:p8L0nBpHJJRKNhu* | *all1440* | N/A | 5.65 |
| Nitrogenase molybdenum-iron protein alpha chain | *nifD* | *ENSB:SBhFQ-pgKfU865y* | *all1454* | N/A | 4.13 |
| Nitrogenase iron protein 1 | *nifH1* | *ENSB:-UiwXBMw740um_c* | *all1455* | N/A | 3.19 |
| Nitrogen fixation protein | *nifU* | *ENSB:zD7Me4dVDd5AIk-* | *all1456* | N/A | 2.11 |
| Nitrogenase cofactor synthesis protein NifS | *nifS* | *ENSB:cwZOUWtsHDDQELT* | *all1457* | N/A | 2.49 |
| Homocitrate synthase | *nifV1* | *ENSB:xnTLVRDJMQvACq9* | *alr1407* | N/A | 4.53 |
| Nitrogen fixation protein | *nifB* | *ENSB:oKdYkJwJmedayE7* | *all1517* | N/A | 2.95 |
| Protein HesB, heterocyst | *hesB* | *ENSB:0W1BDOVJav4vemq* | *all1431* | N/A | 4.23 |
| Protein HesA, heterocyst | *hesA* | *ENSB:Byut0LlNHsEQVLj* | *all1432* | 1.79 | 5.27 |
| Periplasmic [NiFeSe] hydrogenase large subunit | *hupL* | *ENSB:q7HshiWY1tEwz49* | *all0687* | N/A | 3.49 |
| Periplasmic [NiFe] hydrogenase small subunit | *hupS* | *ENSB:YybVoSB-3p1afrt* | *all0688* | N/A | 2.23 |

| | | | | | |
|---|---|---|---|---|---|
| Ferredoxin | *fdxB* | *ENSB:g-hu8fN-yJ9QOkL* | *asr2513* | N/A | 2.91 |
| Ferredoxin, heterocyst | *fdxH* | *ENSB:pgoFbyUO25Y11KZ* | *all1430* | N/A | 3.25 |
| Diflavin flavoprotein flv1B | *flv1B* | *ENSB:egKkM6AIKDL3lV3* | *all0177* | N/A | 1.02 |
| Diflavin flavoprotein flv3B | *flv3B* | *ENSB:gyAx2_YtlGWpP0r* | *all0178* | N/A | 1.32 |
| **Photosynthesis and respiration** | | | | | |
| NTD-OCP like protein | N/A | *ENSB:YZqer7AeouGwj-a* | *all4941* | 1.38 | N/A |
| Homologue of the N-terminal domain of OCP | N/A | *ENSB:Pjo4r6jSQqj9ZKS* | *alr4783* | -1.77 | -2 |
| Homolog of the C-terminal domain of the OCP | N/A | *ENSB:-Skx2hMko_CWcds* | *all4940* | -1.03 | N/A |
| Phycoerythrocyanin subunit beta | *pecB* | *ENSB:DlJg8tfpJb3Pc02* | *alr0523* | -2.57 | -1.92 |
| Phycoerythrocyanin alpha chain | *pecA* | *ENSB:DkyiMvYmj7wtDsq* | *alr0524* | -2.42 | -1.89 |
| Phycobilisome 34.5 kDa linker polypeptide, phycoerythrocyanin-associated, rod | *pecC* | *ENSB:4P_EwWN8WSSWyss* | *alr0525* | -2.75 | -2.03 |

| | | | | | |
|---|---|---|---|---|---|
| Bilin biosynthesis protein PecE | *pecE* | *ENSB:QZBFZRurZXLurvC* | *alr0526* | -3.16 | -2.31 |
| Bilin biosynthesis protein PecF | *pecF* | *ENSB:zGBsC3D8Gtj2ymD* | *alr0527* | -1.94 | -2.2 |
| Photosystem II protein D1 | *psbA 3* | *ENSB:9agEV78l86gFA_V* | *alr4592* | 1.06 | -1.28 |
| Photosystem II protein D1 | *psbA 1* | *ENSB:yJKfo7_EpcR1xJq* | *alr4866* | N/A | N/A |
| Photosystem II CP43 reaction center protein homologue | *isiA2* | *ENSB:n2bv19CnTlJ6ejC* | *all4002* | N/A | 1.89 |
| Photosystem II CP43 reaction center protein homologue | *isiA3* | *ENSB:72J4WLJgrkah1ix* | *all4003* | N/A | 1.76 |
| Light-independent protochlorophyllide reductase iron-sulfur ATP-binding protein | *chlL* | *ENSB:O1DJvDa30jvN8hW* | *all5078* | -1.54 | N/A |
| Oxygen-independent coproporphyrinogen III oxidase | *hemN* | *ENSB:Klj8zn66_ssKF-m* | *alr3126* | N/A | N/A |
| Ferredoxin-1 | *petF* | *ENSB:cn6HVgj9et9zpgG* | *all4148* | N/A | N/A |
| Flavodoxin | *isiB* | *ENSB:vCA6KyCuxG-PG7-* | *alr2405* | N/A | 1.8 |
| Diflavin flavoprotein Flv2 | *flv2* | *ENSB:txsyY6CJmI0pCJY* | *all4444* | N/A | N/A |

| | | | | | |
|---|---|---|---|---|---|
| Diflavin flavoprotein Flv4 | *flv4* | *ENSB:dd_Ejl4qMz5TVdO* | *all4446* | N/A | N/A |
| Cytochrome c oxidase subunit 1 | *coxA2* | *ENSB:6VxYhoGYWmvmjDB* | *alr2515* | N/A | 3.14 |
| Putative cytochrome c oxidase subunit 3 | *coxC2* | *ENSB:Oyzg-naQZMZT0Ld* | *alr2516* | N/A | 4.16 |
| Cytochrome c oxidase subunit 1 | *coxA3* | *ENSB:OjIeYMGIKj96usn* | *alr2732* | N/A | 3.25 |
| Oxidoreductase | N/A | *ENSB:_N-AKaQFkqo5q49* | *all5345* | N/A | 3.06 |
| Pentapeptide repeat protein | N/A | *ENSB:U7IgWWVcVHHMram* | *alr5209* | N/A | N/A |
| Two-component response regulator | N/A | *ENSB:wQizH64rRL3JSms* | *alr0072* | -2.1 | -1.99 |
| **C-metabolism** | | | | | |
| Sbta bicarbonate Na+ symporter | *sbtA* | *ENSB:Z3aL43jCkgjvSJl* | *all2134* | N/A | N/A |
| Bicarbonate transport system permease protein | *cmpB* | *ENSB:Q5MPpHiTaAnt52i* | *alr2878* | N/A | N/A |
| Sucrose synthase | *susB* | *ENSB:49m5v0JcFmdYuMU* | *all1059* | -2.18 | -2.06 |
| Alpha,alpha-trehalase | *treH* | *ENSB:lAnWugFkoPK758N* | *all0166* | -1.79 | -1.94 |

| | | | | | |
|---|---|---|---|---|---|
| Malto-oligosyltrehalose trehalohydrolase | *treZ* | *ENSB:wYIPo5aOxCDd1v_* | *all0168* | -2.27 | -1.98 |
| Transketolase | *tkt_2* | *ENSB:bvaFhPk4hqCTUJt* | *all4052* | -2.4 | -2.21 |
| Alpha-glucanotransferase | N/A | *ENSB:8vPod86j-yWJPGD* | *all0875* | -3.46 | -3.96 |
| 6-Phosphofructokinase | *pfkA1* | *ENSB:STpcob1Kje9jW0R* | *all7335* | N/A | N/A |
| Galactosyltransferase | N/A | *ENSB:F5rC4qxkMvFfiwq* | *all2037* | N/A | 1.91 |
| **Metals** | | | | | |
| Cation-efflux system membrane protein | N/A | *ENSB:jOF_mylngWua0Ez* | *all2845* | N/A | N/A |
| Similar to Na+/H+ antiporter | N/A | *ENSB:9_uVQJPEWe0wqUI* | *all4832* | N/A | -1.27 |
| ABC transporter, ATP-binding protein | *mntA* | *ENSB:PmuYV0bqwlKOUDW* | *all3575* | N/A | -1.13 |
| **Phosphor-metabolism** | | | | | |
| Phosphonate ABC transporter permease | N/A | *ENSB:m4s0rAAffSgr4y7* | *all8088* | N/A | N/A |
| Phosphonate ABC transporter, | N/A | *ENSB:PbMDqwXP9BtwdTI* | *all8089* | N/A | N/A |

| | | | | | |
|---|---|---|---|---|---|
| phosphate-binding protein | | | | | |
| ATP-binding component | *phnL* | *ENSB:LSu_SJZWdPPP-RV* | *all2217* | N/A | N/A |
| Phosphonate ABC transporter, ATP-binding component | *phnK* | *ENSB:L4cUHY-SreLDCy2* | *all2218* | N/A | N/A |
| Phosphonate ABC transporter permease protein | *phnE* | *ENSB:e6bTBbIW4RXbv2I* | *all2227* | N/A | N/A |
| ABC tranporter, phosphate-binding protein | *phnD* | *ENSB:eQSxkgHwKX25u-K* | *all2228* | N/A | N/A |
| ABC tranporter, ATP-binding component | *phnC1* | *ENSB:bURWmyKBm4Qc7CU* | *all2230* | N/A | N/A |
| Phosphodiesterase, alkaline phosphatase D | *phoD* | *ENSB:hgDoJVzVX8lePKi* | *alr2234* | N/A | N/A |
| ABC transporter, periplasmic phosphate-binding protein | N/A | *ENSB:ANyliMgnrcBMIfb* | *alr1094* | N/A | 2.58 |
| **Transcription Factors** | | | | | |
| PacR | *rbcR* | *ENSB:3znthW3skDOmleL* | *all3953* | N/A | N/A |

| | | | | | |
|---|---|---|---|---|---|
| Nitrogen-responsive regulatory protein | *ntcA* | *ENSB:wHiGICXo9Og7Xlz* | *alr4392* | N/A | N/A |
| Cell wall-binding protein Fur | *furA* | *ENSB:D3Qm5IOjvV5LGGN* | *all1691* | N/A | N/A |
| RNA polymerase sigma-subunit | *sigC* | *ENSB:WGvPMBCl6Hl3Gwg* | *all1692* | N/A | 2.56 |
| AraC type transcriptional regulator | N/A | *ENSB:f_q6Xin8pCUNvPv* | *all2035* | N/A | 3.02 |
